# Mechanisms Underlying WNT-mediated Priming of Human Embryonic Stem Cells

**DOI:** 10.1101/2021.05.01.442278

**Authors:** Anna Yoney, Lu Bai, Ali H. Brivanlou, Eric D. Siggia

## Abstract

Embryogenesis is guided by a limited set of signaling pathways dynamically expressed in different places. How a context dependent signaling response is generated has been a central question of developmental biology, which can now be addressed with *in vitro* models of human embryos that are derived from embryonic stem cells (hESCs) called gastruloids. Our previous work demonstrated that during early self-organization of gastruloids, cells chronicle signaling hierarchy. Only cells that have been exposed (primed) by WNT signaling can respond to subsequent Activin exposure and differentiate to mesendodermal (ME) fates. Here, we show that WNT priming does not alter SMAD2 binding nor its chromatin opening, but rather, acts by inducing the expression of the SMAD2 co-factor, EOMES. Expression of EOMES is sufficient to replace WNT upstream of Activin-mediated ME differentiation, thus unveiling the mechanistic basis for priming and cellular memory in early development.

## Introduction

Understanding the transition from pluripotency towards differentiation of human embryonic stem cells (hESCs) is a critical step in elucidating specific aspects of human development. The Activin/Nodal signaling pathway, mediated by transcription factors SMAD2/3, plays a dual role in this process. On one hand, nuclear SMAD2/3 promotes the expression of pluripotency factors *OCT4* and *NANOG* and is essential for hESC pluripotency maintenance (Beattie et al., 2005; James et al., 2005; Vallier et al., 2005; Xiao et al., 2006); on the other hand, in combination with Wnt/β-Catenin signaling, SMAD2/3 can induce hESC differentiation into primitive streak derivatives (Loh et al., 2014; Singh et al., 2012; Sumi et al., 2008), including human organizer tissue (Martyn et al., 2018).

Classical work in vertebrate embryos, originally performed in *Xenopus* embryos and subsequently in fish and mouse, emphasized the importance of Activin/Nodal signaling pathway acting as a morphogen to induce and pattern mesendodermal (ME) lineages including the organizer/primitive streak (Dunn et al., 2004; Green and Smith, 1990; Wilson and Melton, 1994). The idea that Activin acts as a morphogen has consequently been adopted by *in vitro* differentiation studies of hESCs: low levels are required to maintain pluripotency, and moderate to high levels, have been assumed to push the embryonic cells toward ME differentiation. Previous work demonstrated that hESCs can maintain pluripotency over a wide range of Activin concentrations and that Wnt signaling is required for Activin dose-dependent differentiation (Funa et al., 2015; Martyn et al., 2018; Singh et al., 2012; Yoney et al., 2018). The mechanism by which WNT alters the cellular response to different levels of Activin/Nodal signaling, and more generally, how these two signaling pathways synergize to induce mesendoderm (ME) differentiation is not fully understood.

Several mechanisms have been proposed as to how WNT and Activin signaling interact to drive ME differentiation in ESCs. For example, it was proposed that the WNT effector, β-CATENIN, physically interacts with SMAD2/3 to direct its binding and activation of target genes (Funa et al., 2015; Wang et al., 2017); alternatively, it was suggested that β-CATENIN may not directly interact with SMAD2/3, but that it activates transcription synergistically by promoting different transcriptional activation steps (Estarás et al., 2015); β-CATENIN binding may also help SMAD2 overcome a repressor complex, TAZ/YAP/TEADs, at ME specific genes and thus enable their activation (Beyer et al., 2013; Estarás et al., 2017).

A common theme of these mechanistic models is that they are all based on *simultaneous* WNT and Activin stimulation and thus involve co-binding of β-CATENIN and SMAD2/3 to the target ME genes. In contrast, our previous study revealed that the coordination of WNT and Activin/Nodal involves a signaling cascade of WNT acting upstream of Activin: hESCs “primed” with WNT for 24 hours with subsequent exposure to Activin can differentiate into ME cells, while each signaling pathway alone is unable to induce this differentiation (Yoney et al., 2018). Our finding that these two signaling pathways can be *temporally separated* points to a new mechanism that mediates the synergy between WNT and TGFβ signaling in the context of ME differentiation; it also provides a new way to decouple and manipulate the differentiation process.

Using the WNT to Activin stepwise stimulation protocol, we found that WNT priming does not affect the binding or chromatin opening by SMAD2/3, as suggested by previous models. A brief overlap between nuclear β-CATENIN and SMAD2/3 is also not sufficient for hESCs to exit pluripotency, and once hESCs have been primed, nuclear β-CATENIN is dispensable for ME differentiation. Instead, we find that differentiation critically depends on the induction of the transcription factor EOMES during WNT priming, a key regulator of ME differentiation across vertebrates (Arnold et al., 2008; Brown et al., 2011; Bruce et al., 2003; Ryan et al., 1996). Exogenously expressed EOMES can completely replace WNT signaling and drive ME differentiation together with SMAD2/3. Therefore, this study reveals that the primary function of WNT signaling during ME differentiation is not to globally interact with SMAD2/3 but rather to induce EOMES expression.

## Results

### WNT functions temporally up-stream of Activin to specify mesendoderm

We have previously shown that WNT and Activin signaling can be presented in temporal succession to induce ME from pluripotent hESCs (Yoney et al., 2018). 24 h of WNT followed by 24 h of Activin stimulation leads to significant reduction in the expression of the pluripotency marker *SOX2* and strong induction of primitive streak and ME marker genes, *TBXT/BRA* and *EOMES*, and anterior ME marker, *GSC* (Figure 1A– C and Figure S1A, B). In a typical experiment, we observed a high number of cells with ME marker expression. From three independent replicates of the WNT/ACT condition we observed 94%, 99%, and 88% of cells with at least one of the ME makers (BRA, EOMES, or GSC) expressed at a level greater than 3 standard deviations above the average background level in pluripotency (–/ACT). Although EOMES and GSC expression are highly correlated in ME cells, both markers show low correlation with BRA (Figure S1C). These differences in correlation likely reflect the differences in the dynamics of gene expression as BRA is induced in the primitive streak/ME but is turned off as cells further differentiate towards anterior ME and definitive endoderm (DE) (Gu et al., 2004; McLean et al., 2007; Teo et al., 2011).

**Figure 1:**
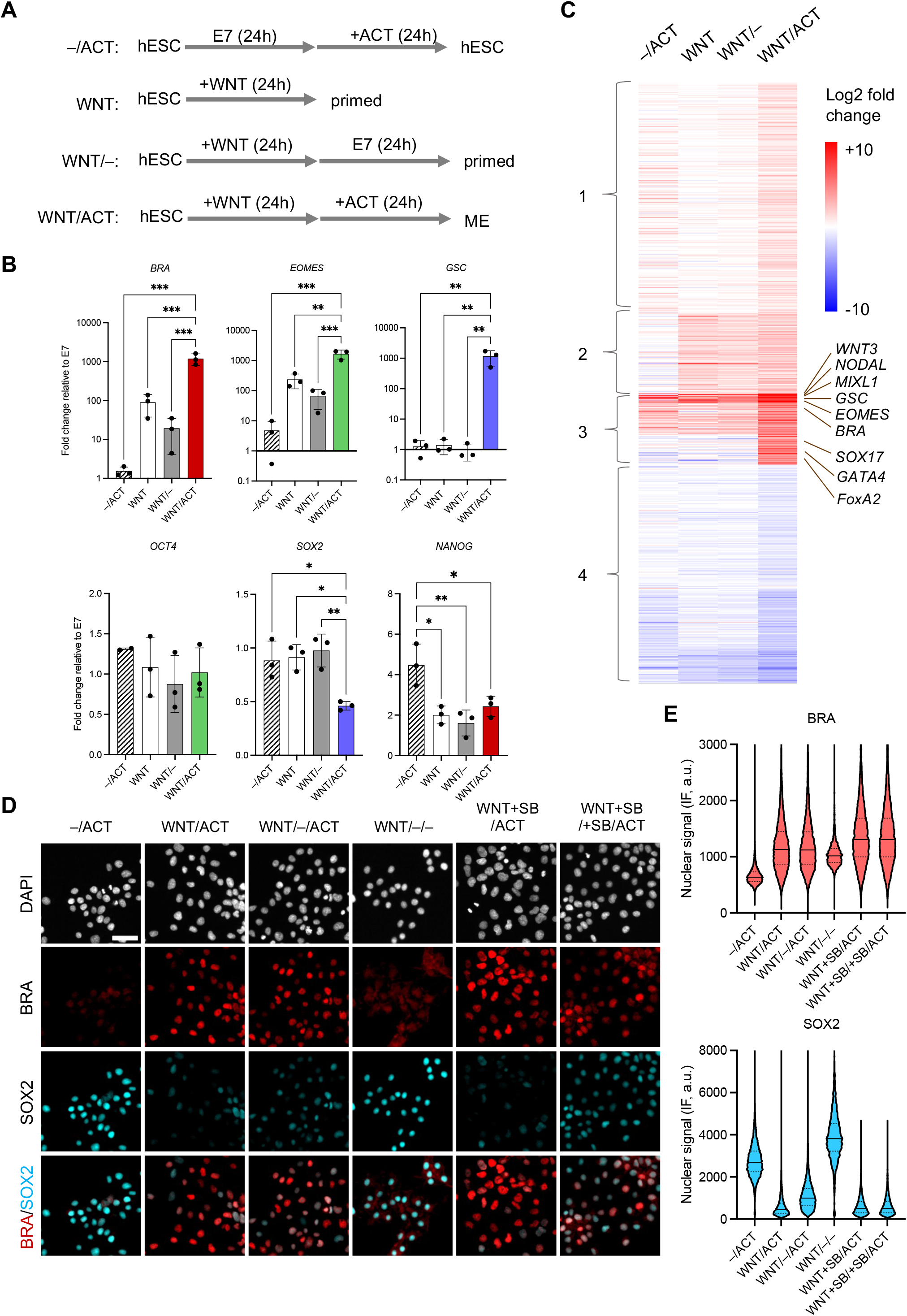
WNT priming and memory in hESCs. **A)** Experimental conditions used to induce a Wnt-primed or mesendoderm state (ME). Cells were treated with 100 ng/mL Wnt3A (WNT) and 10 ng/mL Activin A (ACT), either alone or sequentially, in E7 medium. **B)** Expression of ME genes (*BRA, EOMES, GSC*) and pluripotent genes (*NANOG, OCT4, SOX2*) measured by RT-PCR in cells treated with the experimental conditions defined in (A). Data points represent the mean fold change relative to E7 for independent biological replicates. Bars and error bars represent the mean ± S.D., respectively, across *n* = 3 biological replicates. Differences between means that were determined to be significant are shown (*: p-value < 0.05, **: < 0.01, ***: < 0.001, ANOVA). **C)** *k*-means clustering of RNA-seq data measured in 4 conditions (columns) revealed 4 clusters that could be annotated as follows: genes for which the expression increases in all conditions in which Activin is present (cluster 1, *n* = 415), all conditions in which WNT is present (cluster 2, *n* = 224), or further activated by Wnt and Activin (higher induction in WNT/ACT than WNT or ACT alone, cluster 3, *n* = 137). The fourth cluster represents genes that are repressed by the WNT/ACT condition (cluster 4, *n* = 435). The intensities represent log2 fold change of expression relative to the E7 condition. Only genes showing a significant change (adjusted p-value < 0.01) in at least one condition relative to E7 were included in the heatmap and clustering procedure. **D)** Memory of the WNT-primed state was tested by incubating cells for an additional day in E7 base medium in the presence or absence of SB431542 (SB, 10 μM) prior to presenting ACTIVIN (10 ng/mL). SB was used to eliminate Activin signaling during the first 48 h. After 24 h of Activin stimulation, cells were fixed and analyzed by immunofluorescence (IF) for BRA and SOX2 expression. Scale bar, 50 μm. **E)** Violin plots of the nuclear IF signal quantified in single cells (n > 5,000 cells per condition) from the experiment shown in B.

The results following WNT/ACT contrasts with 24 h of WNT stimulation where the expression of *OCT4* and *SOX2* is maintained at pluripotency levels, *GSC* remains low, and *EOMES* and *BRA* are only mildly activated (Figure 1B and Figure S1A, B). *NANOG* expression is reduced, consistent with the removal of Activin during WNT priming (Vallier et al., 2009). When cells were primed with WNT and then cultured in E7 base medium (Chen et al., 2011) without Wnt or Activin for an additional 24 h, *EOMES* and *BRA* transcript levels start to drop towards their basal levels in E7 (Figure 1B). These observations suggest that 24 h of WNT signaling does not cause a stable fate change in hESCs. When Activin was applied prior to WNT, ME gene expression remains low, strongly suggesting that ME differentiation does not simply require both signals in close succession but also in a particular temporal order (Figure S1A, B).

As shown in Figure S2, similar results were obtained with the small molecule, CHIR99021 (CHIR), which activates the canonical WNT signaling pathway in a cell autonomous manner by blocking β-CATENIN degradation (Kunisada et al., 2012; Naujok et al., 2014). Taken together, these results demonstrate that WNT priming through β-CATENIN is required to switch the output of Activin signaling from pluripotency maintenance to ME speciation.

### WNT priming regulates the expression of a small subset of SMAD2/3-target genes

The experiments presented above confirm our original observation that hESCs have a differential transcriptional response to Activin alone, versus Activin stimulation following WNT priming. To begin to decipher the mechanisms underlying this difference, we performed bulk RNA-sequencing (RNA-seq) under a set of WNT and Activin conditions (Figure 1A). We identified 1212 genes that showed a significant change in expression in at least one of the conditions relative to 24 h in E7 medium (*n*=2 biological replicates, adjusted P-value < 0.01). These genes fall into four major clusters that are induced by Activin only (1), WNT-only (2), WNT together with Activin (3), or repressed under at least one condition (4) (Figure 1C and Table S1).

Among all differentially expressed genes, 189 had significant changes after 24h of WNT presentation. Genes that were induced by WNT, including *EOMES, BRA*, and *NODAL*, partially reversed to their basal level after a subsequent 24 h in E7 medium, which further supports the notion that WNT signaling primes cells rather than causing a stable fate change. The expression of pluripotency makers *NANOG, OCT4, SOX2*, and *KLF4*, either remained constant or was upregulated in response to WNT or Activin, confirming the lack of differentiation with either signal alone. Consistent with broad transcriptional changes associated with differentiation, there were more extensive changes when WNT was followed by Activin (WNT/ACT), including 661 upregulated genes and 383 downregulated genes. In our WNT/ACT condition, cells express markers of anterior ME and early definitive endoderm (DE), including *EOMES, GSC, SOX17, FOXA2*, and *GATA4* (Figure 1C and Table S1). We do not observe expression of key markers of cardiac, paraxial, or lateral plate mesoderm, including *MESP1/2, TBX6, MSGN1/2, HAND1*, or *ISL1*. These results are consistent with previously published *in vivo* and *in vitro* studies, which demonstrated that cells differentiate towards the DE fate via an ME intermediate (Gu et al., 2004; McLean et al., 2007; Teo et al., 2011).

In our previous study we showed that the application of Activin to hESCs induces transient translocation of SMAD2 and SMAD4 into the nucleus, which elicits a transcriptional change in ~3500 genes (Yoney et al., 2018). Many of the differentially expressed genes are activated rapidly following SMAD2/4 translocation and are repressed after its nuclear exit, suggesting that they are direct targets of SMAD2/4. Among these genes, 113 of them show significantly higher mRNA levels in WNT/ACT compared to WNT or ACT only conditions, indicating that these genes respond to the cooperative signaling of WNT and Activin. This group of genes includes *EOMES, BRA, GSC, MIXL1, GATA6 and NODAL*, which agrees with previous reports that β-CATENIN and SMAD2/3 co-regulate expression of genes that are expressed within the primitive streak and ME derivatives (Estarás et al., 2015; Funa et al., 2015). The remaining SMAD2/4 targeting genes identified in our previous study are not sensitive to WNT priming.

In summary, our genome-wide transcriptional analyses support the idea that 24h of WNT stimulation primes pluripotent hESCs for differentiation and that together, WNT and Activin drive cells towards a committed ME fate. We were able to unbiasedly identify a group of Activin/SMAD2 target genes whose transcriptional responses are affected by WNT priming (RNA levels under WNT/ACT are significantly higher than those in WNT/- and -/ACT). In our subsequent analyses, we will compare WNT priming dependent vs independent genes with respect to additional features, including transcription factor (TF) binding sites and chromatin accessibility, to further dissect the mechanism of WNT priming.

### Cells retain a memory of WNT priming

Motivated by the observations of “Wnt memory” in other species ranging from flies to frog (Alexandre et al., 2013; Blythe et al., 2010), we further explored our original hypothesis that hESCs possess the ability to record WNT signals. To demonstrate extended WNT memory, we stimulated cells with WNT for 24 h, followed by one additional day in E7 medium before switching to Activin for 24 h. The small molecule inhibitor SB431542 (SB) was also included to block the cells’ ability to respond to endogenous Activin/Nodal signals and up-regulate ME markers. Consistent with previous observation (Yoney et al., 2018), adding SB during Wnt priming has little effect on the subsequent Activin response (Figure 1D, E). Despite the extended interval between WNT and Activin stimulation, Activin can still induce ME markers and down-regulate SOX2 at the protein level, albeit to a lesser extent (Figure 1D, E and Figure S3). We interpret these results to mean that 1) the WNT effect decays over time, and the cells are not stably committed to the “WNT-primed” state, and 2) the WNT priming effect is still available for at least a day after removal of the signal.

### SMAD2 binds to the same loci with or without WNT priming

We next investigated the mechanism by which WNT priming modifies the cellular response to Activin. We had previously shown that SMAD2 concentration and nuclear residency dynamics are not affected by WNT priming, suggesting that WNT affects SMAD2 activity in a subsequent step (Yoney et al., 2018). Since SMADs have weak DNA specificity and often rely on co-factors to be directed to target genomic regions in a cell-type specific manner (Massagué, 2012), we first considered the possibility that WNT priming alters the target sites of SMAD2/4. Accordingly, we analyzed the SMAD2/3 ChIP-seq data in Kim et al., 2011 (ChIP in hESCs in conditioned media, which is similar to our E7(–)/ACT condition) and in Tsankov et al., 2015 (ChIP in ME cells generated with WNT + ACT treatment for 12h, which is similar to our WNT/ACT condition). The overall SMAD2/3 ChIP profiles are similar in these two conditions.

Several examples are shown in Figure 2A, where the SMAD2 ChIP-seq signals are largely overlapping despite the fact that the mRNA levels of these genes are a few fold or even orders of magnitude higher in the WNT priming followed by Activin condition (WNT/ACT) than in Activin alone condition (–/ACT) (Figure S4). We also quantified the area underneath the SMAD2/3 ChIP-seq peaks from these datasets, and they are highly correlated, both at the genome-wide scale (R ~0.8; Figure 2B), or in the vicinity of genes that are sensitive to WNT priming (R = 0.86; Figure 2C). ChIP data tend to be noisy, and even among the three replica of the ME ChIP data, the average correlation is only ~0.71. We, therefore, conclude that SMAD2 binding measured in these two datasets/conditions is highly similar.

**Figure 2:**
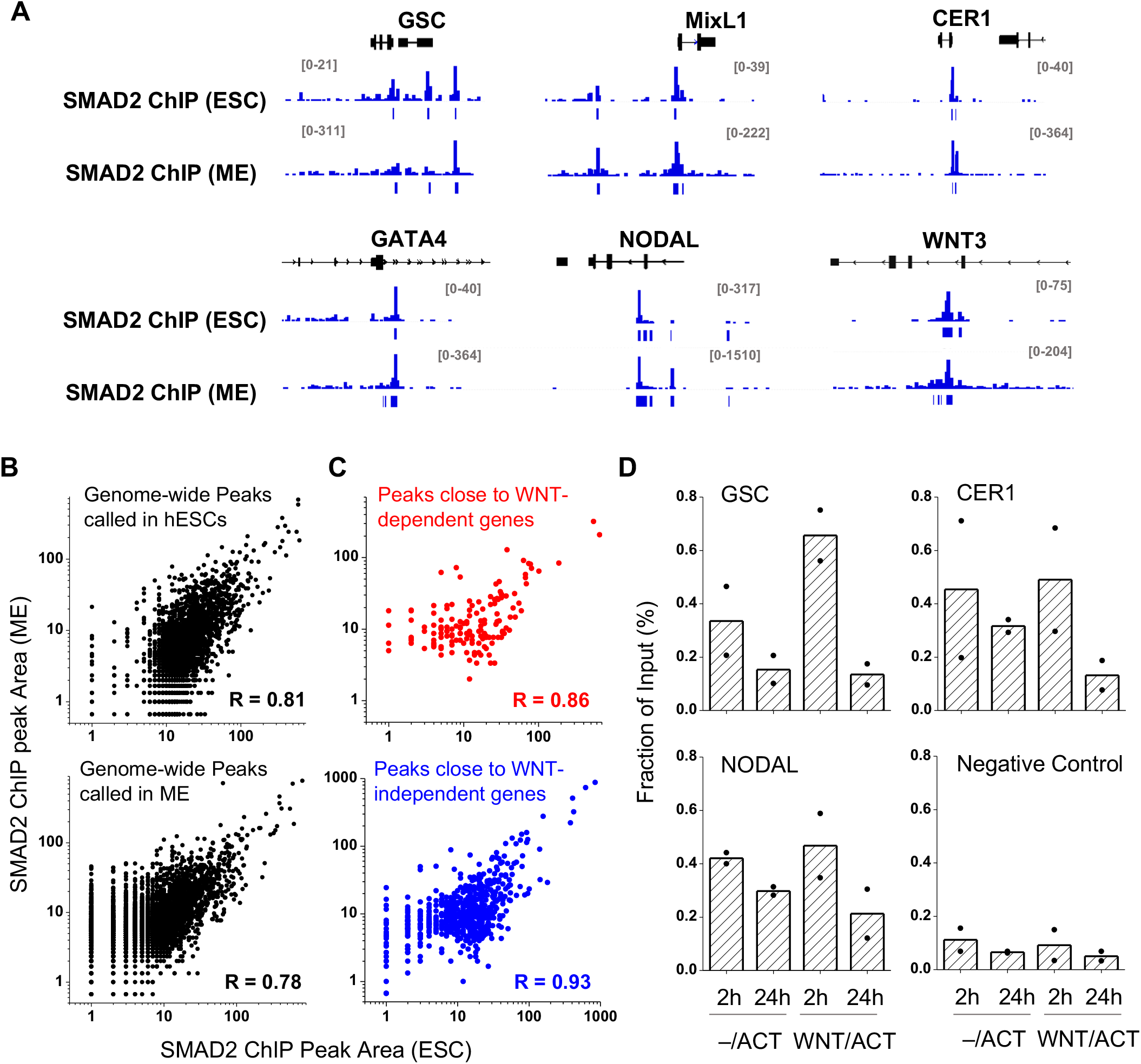
SMAD2 binds the same genomic loci with or without WNT priming. **A)** SMAD2/3 ChIP data from pluripotent hESCs (Kim et al. 2011) or hESC derived ME cells (Tsankov et al. 2015) in the vicinity of *GSC*, *MIXL1, CER1, GATA4, NODAL*, and *WNT3* genes. Each region is ~20 kb in size. These genes are significantly upregulated in ME cells, but the SMAD2/3 binding patterns remain largely unchanged. **B)** Genome-wide SMAD2 ChIP signals in hESCs vs. ME derived from the same datasets as in (A). The ChIP signals were qualified using the area underneath the ChIP peaks. Using either peaks called in hESCs or ME cells, the levels of SMAD2 binding are similar in these two cell types. R, Pearson correlation. **C)** Same as in (B) but restricted to SMAD2 ChIP peaks near target genes (peaks within 10kb upstream of TSS and 10kb downstream of TES) that are either affected (upper panel) or unaffected (lower panel) by WNT priming. **D)** SMAD2 ChIP signals over previously known SMAD2 binding sites near *GSC*, *CER1*, and *NODAL* genes, as well as a negative control region that does not show SMAD2 binding in the genome-wide datasets. The ChIP measurements were performed at 2 or 24 h after Activin (10 ng/mL) application with or without Wnt priming. The ChIP signal was normalized by the total input. Bars represents the mean for n = 2 biological replicates, and individual measurements are shown in the plot.

Since the published SMAD2/3 ChIP data were performed in cell lines and conditions that are not identical to ours, we selected a few SMAD2 binding regions near genes that respond to WNT priming (*GSC*, *CER1*, and *NODAL*) and carried out ChIP-qPCR measurements at two different time points during Activin treatment with or without WNT priming (Figure 2D). For all genes tested, we observed an increase in SMAD2 binding at 2 h relative to 24 h following Activin presentation, consistent with the adaptive change of the SMAD2 nuclear concentration over time (Yoney et al., 2018). However, the level of binding is comparable in the presence or absence of WNT priming. This analysis indicates that higher SMAD2 nuclear concentration promotes SMAD2 binding at target genes. However, SMAD2 binding *per se* is not correlated with transcriptional output and corresponding changes in cell fate.

### SMAD2 leads to a comparable level of chromatin opening with or without WNT priming

Given that SMAD2 can bind to the same genes with or without WNT priming but activates them to a different extent, it is possible that SMAD2 may have different activation potentials with or without WNT priming. Because TF activation is often accompanied by local chromatin opening, we carried out ATAC-seq to probe the changes in chromatin accessibility. The ATAC-seq experiments were performed in the same conditions described above for our bulk RNA-seq analysis. The ATAC-seq profiles in all conditions are very similar showing that there is no global rearrangement of open chromatin regions in the transition from pluripotency to ME differentiation (example traces in Figure 3A and global analysis in Figure S5A, B). Furthermore, a vast majority of genes contain ATAC-seq peaks near their TSS and within transcript regions, regardless of their transcriptional status.

**Figure 3:**
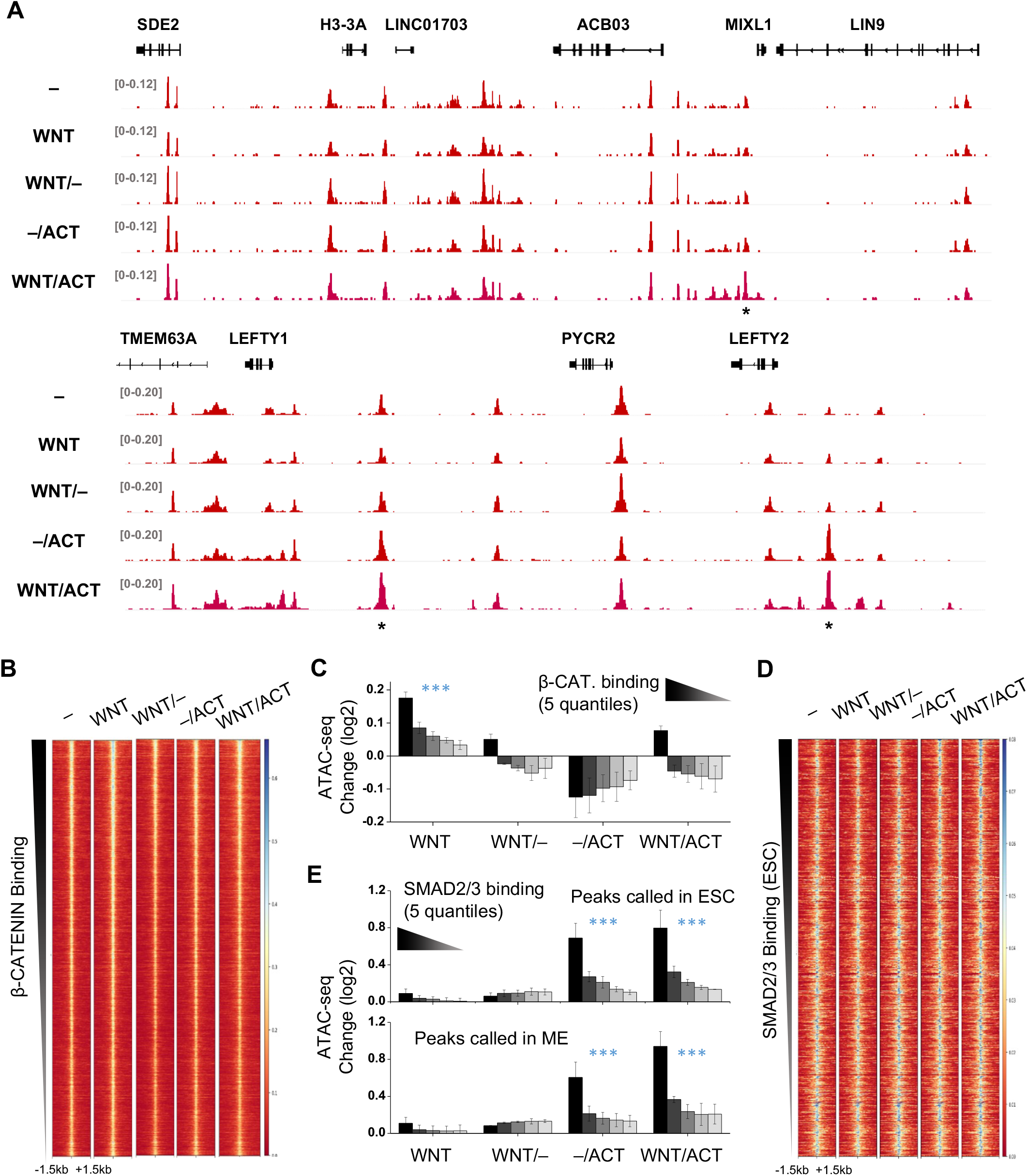
SMAD2 shows similar chromatin opening activity with or without WNT priming. **A)** ATAC-seq data over two example regions measured in five conditions as indicated. The overall pattern looks identical except for a few peaks marked with (*). Cells without treatment were cultured in E7(–). **B)** ATAC-seq signals centered at β-CATENIN binding sites (n = 17809) in five conditions as in (A). β-CATENIN binding sites in WNT-treated ESCs were derived from published ChIP-seq data (Estarás, Benner, and Jones 2015). Rows represent genomic regions sorted by the β-CATENIN ChIP peak intensities. **C)** Fold change of the ATAC-seq signals over the β-CATENIN binding sites in four conditions relative to E7. The five bars in each condition represent 5 quantiles of β-Catenin binding strength. The ATAC-seq signals are significantly enhanced under the Wnt condition (p-value < 0.001 for all five quantiles, paired t test). **D**) Same as in (B) except that ATAC-seq signals are centered at SMAD2/3 binding sites (n = 4032) based on ChIP data from pluripotent hESCs (Kim et al. 2011). **E)** Same as in (C) except that the five bars represent 5 quantiles of SMAD2/3 binding strength (based on ChIP data in either pluripotent hESCs or ME cells). The ATAC-seq signals are significantly enhanced under the –/ACT and WNT/ACT condition (p-value < 0.001 for all five quantiles, paired t test).

Despite the global similarities, there are quantitative changes in the ATAC-seq signals over specific TF binding sites. Two examples of ATAC-seq signal change near *MIXL1, LEFTY1* and *LEFTY2* are shown in Figure 3A (peaks marked with *). These dynamic ATAC-seq peaks overlap with the binding sites of several TFs, including β-CATENIN, SMAD2, and EOMES (Figure S5C). For β-CATENIN, the ATAC-seq data clearly show that globally its binding sites are accessible in E7 medium prior to WNT stimulation (Figure 3B). After 24 h WNT treatment, β-CATENIN associates with these pre-opened chromatin regions and leads to mild but statistically significant enhancement in ATAC-seq signals (Figure 3C). The level of enhancement correlates with the strength of β-CATENIN binding determined from ChIP-seq peak area (Estarás et al., 2015). Interestingly, such enhancement is reduced when cells are moved back to E7 medium following WNT priming (WNT/–), indicating that β-CATENIN at least partially dissociates from chromatin after elimination of WNT signals. In addition, the ATAC-seq peaks drop to the same level when cells are primed with WNT and then stimulated with Activin (WNT/ACT) as in the WNT/– condition, suggesting that the presence of nuclear SMAD2 does not retain β-CATENIN on chromatin (Figure 3C).

We performed the same analysis using SMAD2 binding sites (Kim et al., 2011; Tsankov et al., 2015). The ATAC-seq signals over the SMAD2 binding sites are essentially identical in E7 base medium and WNT conditions, indicating that WNT signaling does not alter the chromatin environment near these sites. In contrast, with both Activin only (–/ACT) and WNT priming followed by Activin treatment (WNT/ACT), ATAC-seq signals are significantly enhanced over SMAD2 binding sites, and the level of enhancement correlates with SMAD2 binding strength (Figure 3D, E and Figure S5D). Therefore, such enhancement likely results from SMAD2 binding and subsequent opening of the local chromatin. Importantly, we observed similar enhancement in –/ACT and WNT/ACT samples over the SMAD2 binding sites, which supports our conclusion that SMAD2/3 binding is similar in these two conditions. Moreover, these data indicate that the activity of chromatin opening by SMAD2 is not dependent on WNT priming.

### Nuclear β-CATENIN is insufficient to drive ME differentiation with SMAD2

We have shown that WNT priming affects neither SMAD2 binding nor its ability to open chromatin. It has been proposed that the main TF that responds to WNT signaling, β-CATENIN, functions as a co-activator of SMAD2, either through direct physical interaction (Funa et al., 2015) or by cooperatively recruiting RNA pol II (Estarás et al., 2015). β-CATENIN indeed tends to bind WNT-primed genes together with SMAD2 (Figure S6A, B). It is thus possible that during Activin treatment after WNT priming, the remaining β-CATENIN nuclear fraction synergizes with newly translocated SMAD2 to drive the expression of key ME genes (Figure 4A).

**Figure 4:**
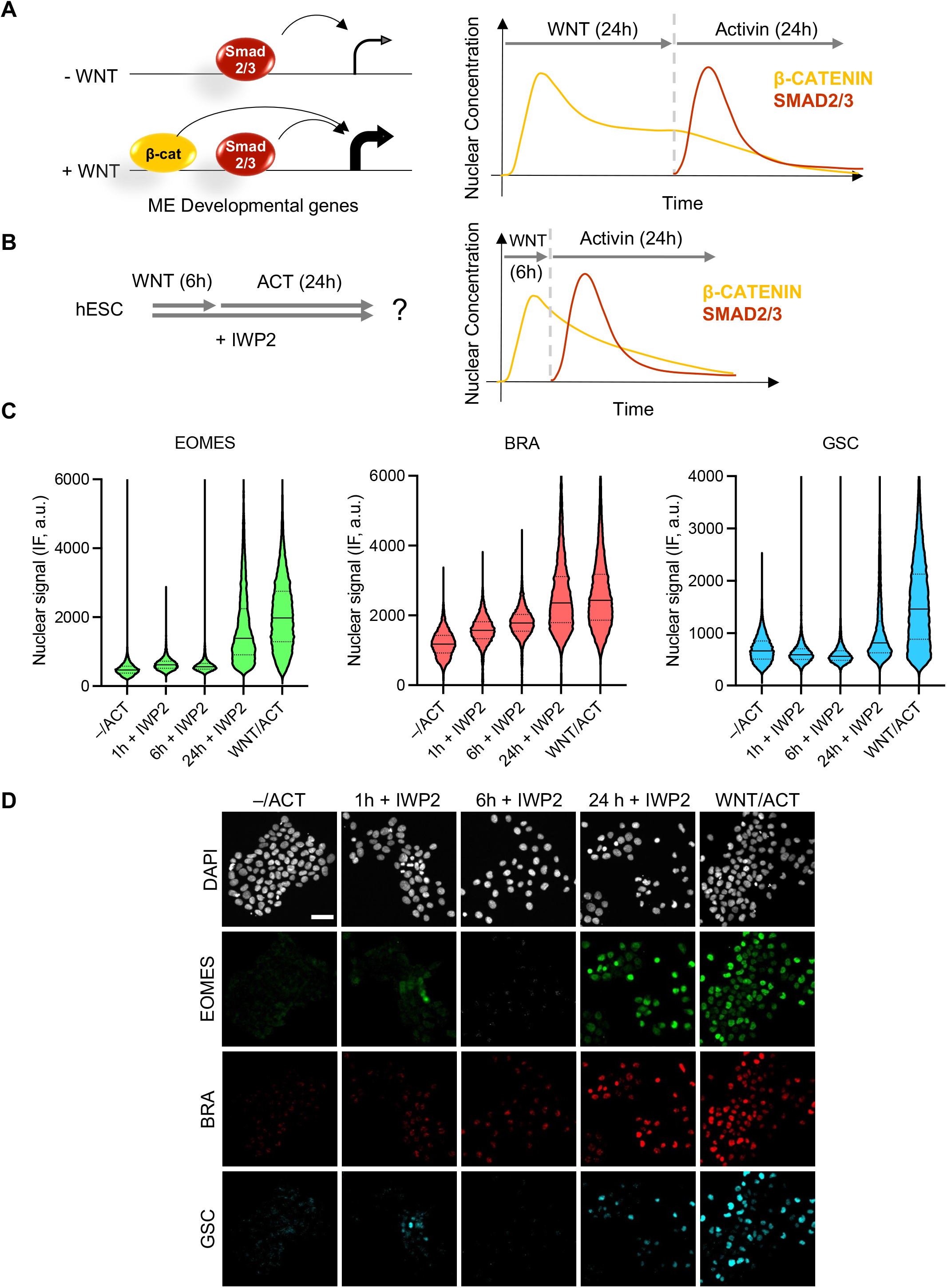
Presence of nuclear β-CATENIN is not sufficient for ME differentiation. **A)** Schematic depicting the hypothesis to be tested that increased ME gene expression requires overlap of β-CATENIN and SMAD2/3. Cartoon showing the nuclear concentration of β-CATENIN and SMAD2 during the WNT/ACT treatment. **B)** Method for testing the hypothesis in (A) using shorter WNT treatment. IWP2, which prevents cells from secreting Wnt ligands, is used in this protocol to ensure that WNT signaling is restricted to the first 6 h. **C)** Short Wnt exposure is not sufficient to prime the cells for ME differentiation. Cells were treated for different amounts of time with WNT (100 ng/mL) prior to adding Activin (10 ng/mL). After 24 h of Activin stimulation, cells were fixed and analyzed by immunofluorescence for EOMES, BRA, and GSC expression. Violin plots of the nuclear IF signal quantified in single cells (n > 10,000 cells per condition). Solid line, median; Dashed lines, upper and lower quartiles. **D)** Example images corresponding to the analysis shown in (C). Scale bar, 50 μm.

Two pieces of evidence argue against this scenario as being the only requirement for ME gene induction. First, β-CATENIN rapidly accumulates in the nucleus after Wnt presentation, reaching the peak concentration at ~6h in hESCs (Massey et al., 2019). If the overlap between nuclear β-CATENIN and SMAD2/4 is the key mechanism of WNT priming, a shorter period of WNT treatment should be sufficient for ME differentiation (Figure 4B). However, 6h or less of WNT priming followed by WNT washout and 24 h of Activin in the presence of an in inhibitor of endogenous WNT secretion (IWP2), is not able to activate ME genes (Figure 4C, D). The fact that WNT priming requires longer time strongly indicates that a product resulting from β-CATENIN activation, instead of β-CATENIN itself, is essential to turn on the ME differentiation program with SMAD2

Second, we tested if the overlap between β-CATENIN and SMAD2 is necessary for ME differentiation. We repeated our WNT/ACT protocol adding the small molecule inhibitor endo-IWR1 at varying times from 0h to 12h after the addition of Activin (Figure 5A). endo-IWR1 promotes the degradation of β-CATENIN through stabilization of the AXIN destruction complex (Chen et al., 2009) and therefore should reduce its overlap with SMAD2 (Figure 5A). Application of this drug together with WNT completely abolishes ME and DE differentiation and maintains pluripotency gene expression (Figure 5B–D). When endo-IWR1 was added immediately after Wnt treatment (0h) together with Activin, the induction of some ME genes was reduced, indicating that a short overlap between β-CATENIN and Activin signaling indeed is important for ME differentiation. However, when the drug was added a few hours (4-12 hours) after the start of the Activin treatment, it no longer blocks ME induction or the downregulation of *SOX2* and *NANOG* (Figure 5B, D). Importantly, marker genes for DE, including *FOXA2, SOX17*, and *GATA4*, were also induced with the later application of endo-IWR1, indicating that hESCs can differentiate into more mature endoderm after the elimination of β-CATENIN (Figure 5C). We note that the expression of all genes tested is stable between 24 and 48 h of Activin treatment except for *BRA*, which is downregulated at 48 vs 24 h (Figure 5B). This is consistent with previous findings that BRA is turned off as cells differentiate from an ME intermediate to more mature DE (D’Amour et al., 2005; McLean et al., 2007). Taken together with experiments presented in Figure 4, we conclude that the presence of β-CATENIN in the nucleus is not sufficient to carry out the ME to DE differentiation program, and after 24 h WNT priming, continuous overlap between β-CATENIN and SMAD2 is also not necessary for ME and early DE differentiation.

**Figure 5:**
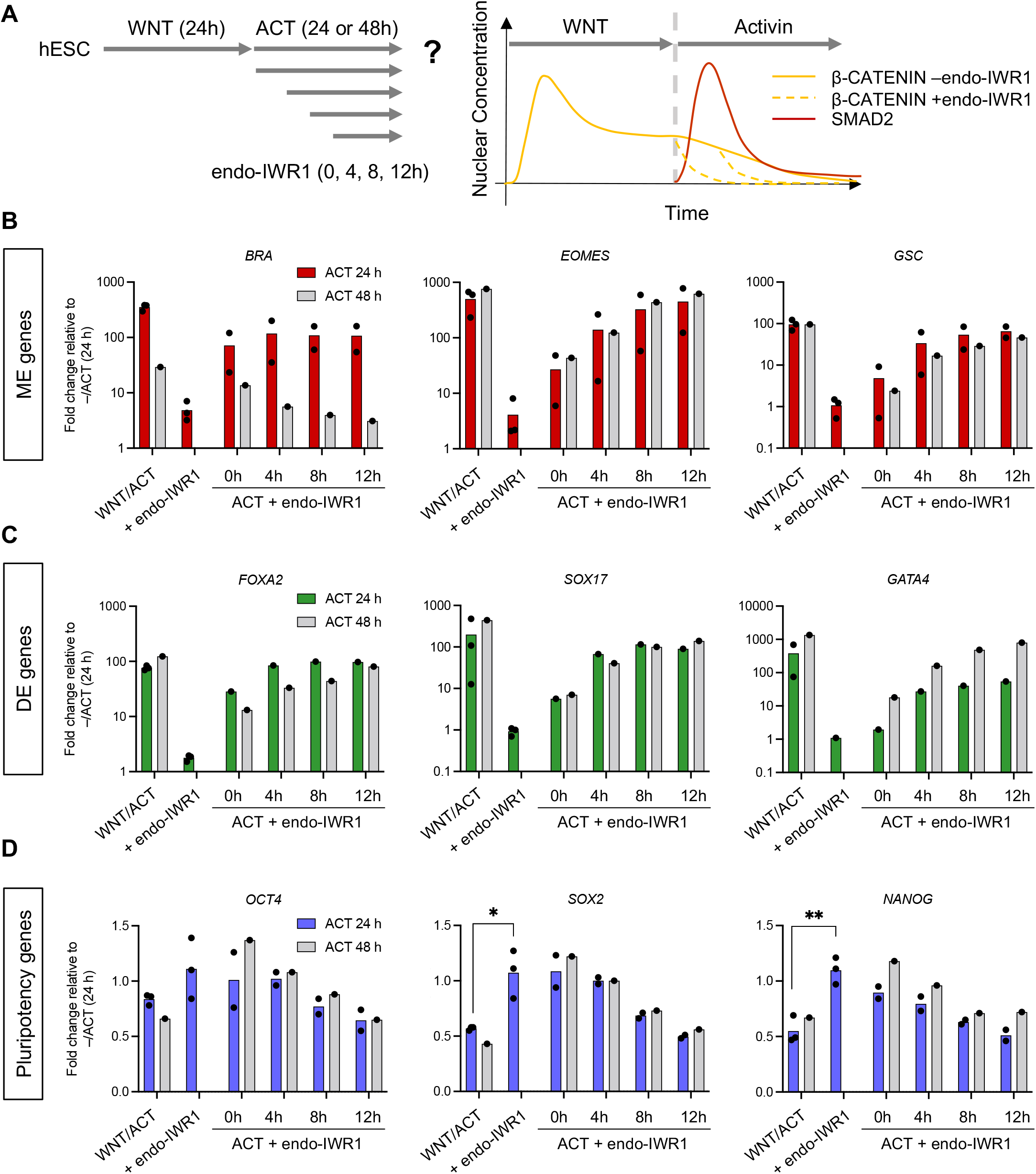
Continuous overlap between β-CATENIN and SMAD2 is not necessary for ME differentiation. **A)** Protocol for testing the effect of different durations of β-Catenin/SMAD2 overlap on ME differentiation. We applied endo-IWR1 (1 μM) with WNT and Activin or at different time points during the Activin phase only to reduce the nuclear concentration of β-CATENIN, thus reducing its overlap with SMAD2. **B–D)** RT-PCR analysis of ME (B), DE (C), or pluripotent genes (D) after 24 h or 48 h of Activin treatment (10 ng/mL) with endo-IWR1 (1 μM) added at different time points. Expression in each sample was normalized to GAPDH and then to the pluripotency levels (–/ACT, 24 h). Individual data points represent biological replicates, and bars indicate the mean. In (D) Student’s t-test was used to compare the mean of WNT/ACT vs the mean of WNT/ACT + endo-IWR1 added throughout the entire protocol (*: p-value < 0.05, **: < 0.01).

### EOMES is a potential effector for WNT priming

Based on the results presented above, we hypothesized that WNT priming generates another TF that functions as an essential SMAD2 co-regulator (Figure 6A). Given our data in Figures 4 and 5, we suspect that this TF should be activated by the end of the 24 h of WNT treatment and should bind in proximity to WNT-primed genes in a WNT/ACT specific manner. Accordingly, we probed the binding signature of this TF by searching for enhanced ATAC-seq peaks that require the combined action of WNT and Activin, e.g. the ATAC-seq peak near the *MIXL1* gene in Figure 3A. We identified 3983 peaks that are significantly enhanced in Wnt priming followed by Activin condition (WNT/ACT) in comparison to both Activin alone (–/ACT) and Wnt alone (WNT/–) (Figure 6B). The probability for enhanced peaks to fall within 20k bp of WNT-primed genes is much higher than for genes that are not primed by WNT, indicating that these ATAC-seq peaks correlate with transcriptional activation and are therefore likely to represent enhancers (Figure 6C).

**Figure 6:**
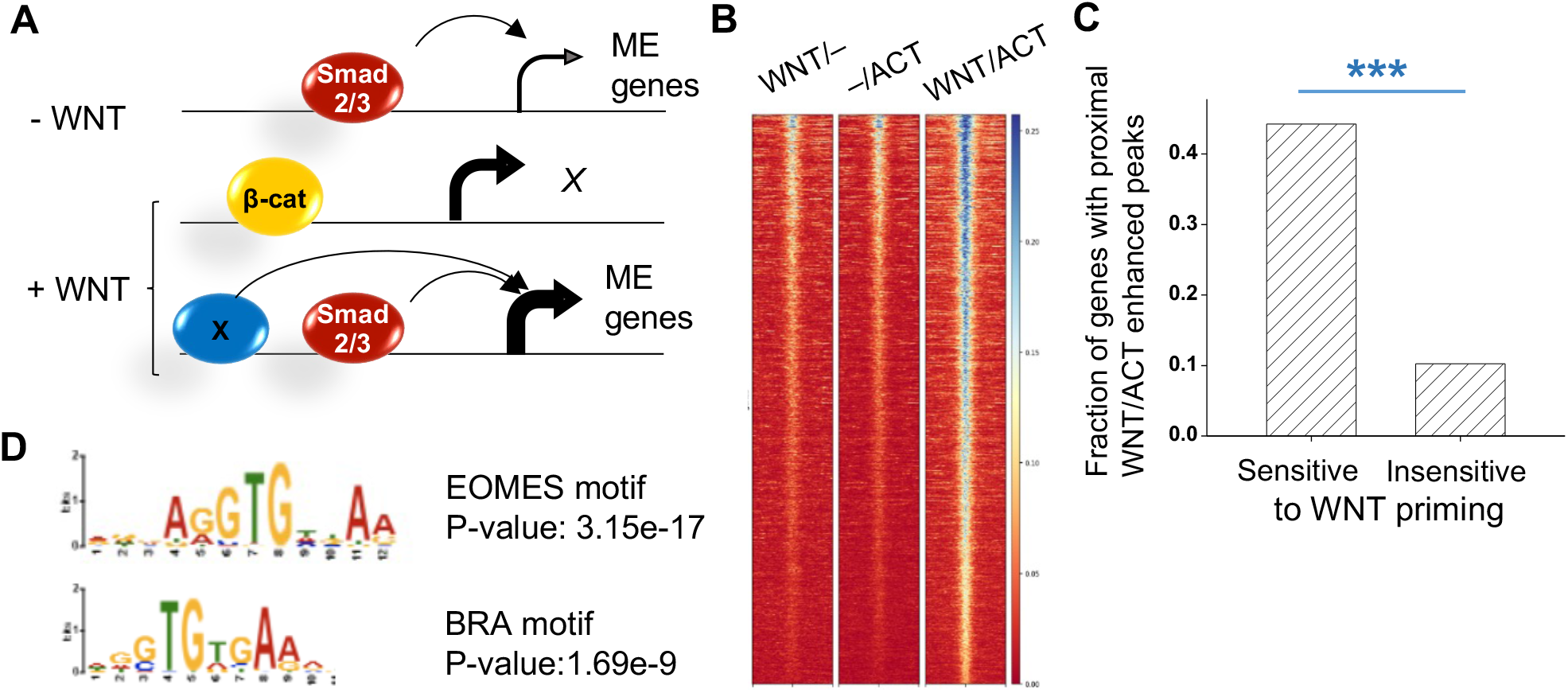
EOMES motif is enriched in WNT/ACT-enhanced ATAC peaks. **A)** Schematic depicting the hypothesis to be tested that increased ME gene expression requires a SMAD2/3 co-activator (X) that is induced by WNT. **B)** Heatmap of the ATAC-seq signals that are enhanced in WNT/ACT condition in comparison to WNT or ACT alone. **C)** Probability of finding these WNT/ACT-enhanced peaks near genes (start of the gene - 20kb to end of the gene + 20kb) that are either sensitive or insensitive to WNT priming. ***: p-value < 10^-4^. **D)** EOMES motif is the most enriched motif, followed by BRACHYURY (BRA) found in the WNT/ACT-enhanced peaks near genes that are WNT primed.

We analyzed motifs that are enriched in the WNT/ACT-enhanced ATAC peaks, both genome-wide and the subset that is proximal to WNT-primed genes. In both cases, we found the EOMES motif to be the most significantly enriched motif (the EOMES motif was detected in ~80% of the sequences) (Figure 6D; full list shown in Table S2). Indeed, ~80% of WNT/ACT-enhanced ATAC-seq peaks overlap with EOMES ChIP peaks, whereas the overlap with β-CATENIN and SMAD2 peaks is much less (Figure S6C). Consistent with our expectation, EOMES is activated by WNT alone and becomes highly expressed with WNT priming followed by Activin (Figure 1B and Table S1). EOMES tends to bind near SMAD2 in the vicinity of Wnt-primed genes (Figure S6D, E), and is an essential TF for differentiation into ME cells and definitive endoderm (DE) (Li et al., 2019; Teo et al., 2011; Tosic et al., 2019). The TF with the second most enriched motif, BRA (Figure 6D), is not essential for DE differentiation *in vitro* (Li et al., 2019; Tosic et al., 2019). Although BRA deficient mice die by embryonic day 10, they do not exhibit substantial defects in the anterior mesendoderm and definitive endoderm but fail to form midline and posterior mesoderm (Wilkinson et al., 1990). The other significantly enriched TF-family motif belongs to the GATA factors. GATA4/6 are key endoderm TFs, but their expression is not upregulated after 24 h WNT (Table S1). Therefore, we conclude that EOMES is the most likely effector of Wnt priming.

### Exogenous EOMES can replace the effect of Wnt

If EOMES is the key factor mediating WNT priming, the prediction is that we can achieve ME differentiation in the absence of WNT by artificially expressing EOMES. To test this hypothesis we generated an hESC line with doxycycline(dox)-inducible *EOMES (TT-EOMES*, Figure S7A) using the piggyBac transposon system (Lacoste et al., 2009). When these cells were treated with dox, morphological changes were induced that were indicative of ME differentiation (Figure S7B). According to previous studies, *EOMES* expression in the absence of Activin drives cells towards cardiac mesoderm fate by activating *MESP1*, and addition of Activin inhibits cardiac differentiation (Ameele et al., 2012). We observed the same trend of *MESP1* expression in our dox-treated TT-EOMES lines in the presence or absence of Activin (Figure S7C). To prevent cells from differentiating towards this alternative path, we added Activin when inducing EOMES with DOX in the protocols below.

We next tested if a pulse of EOMES expression could replace WNT priming to drive ME differentiation. A titration experiment showed that exogenous *EOMES* is induced in a dox concentration- and time-dependent manner (Figure S8A). In particular, 6h treatment with 0.1 μg/mL dox induces EOMES protein expression to a level comparable to the endogenous levels in our WNT/ACT protocol (Figure S8B–C). We induced *TT-EOMES* for 6h with 0.05, 0.1, and 0.2 μg/mL dox in the presence of Activin, washed out dox, and measured gene expression at different time points after dox removal (Figure 7A). *TT-EOMES* expression level decreases rapidly following the removal of dox (Figure 7B). Such transient *TT-EOMES* expression induced by 0.1 and 0.2 μg/mL, but not 0.05 μg/mL dox, was sufficient to drive expression of ME and DE genes, including the endogenous *EOMES*. Except for *BRA*, the levels induced in the TT-EOMES line matched those obtained with WNT/ACT treatment of the unmodified parental cell line (Figure 7C). We noted that ME genes like *EOMES* and *GSC* are activated earlier than *bona fide* DE genes like *SOX17*, indicating that the cells driven by *TT-EOMES* go through a differentiation process similar to that obtained with other *in vitro* DE protocols (McLean et al., 2007).

**Figure 7:**
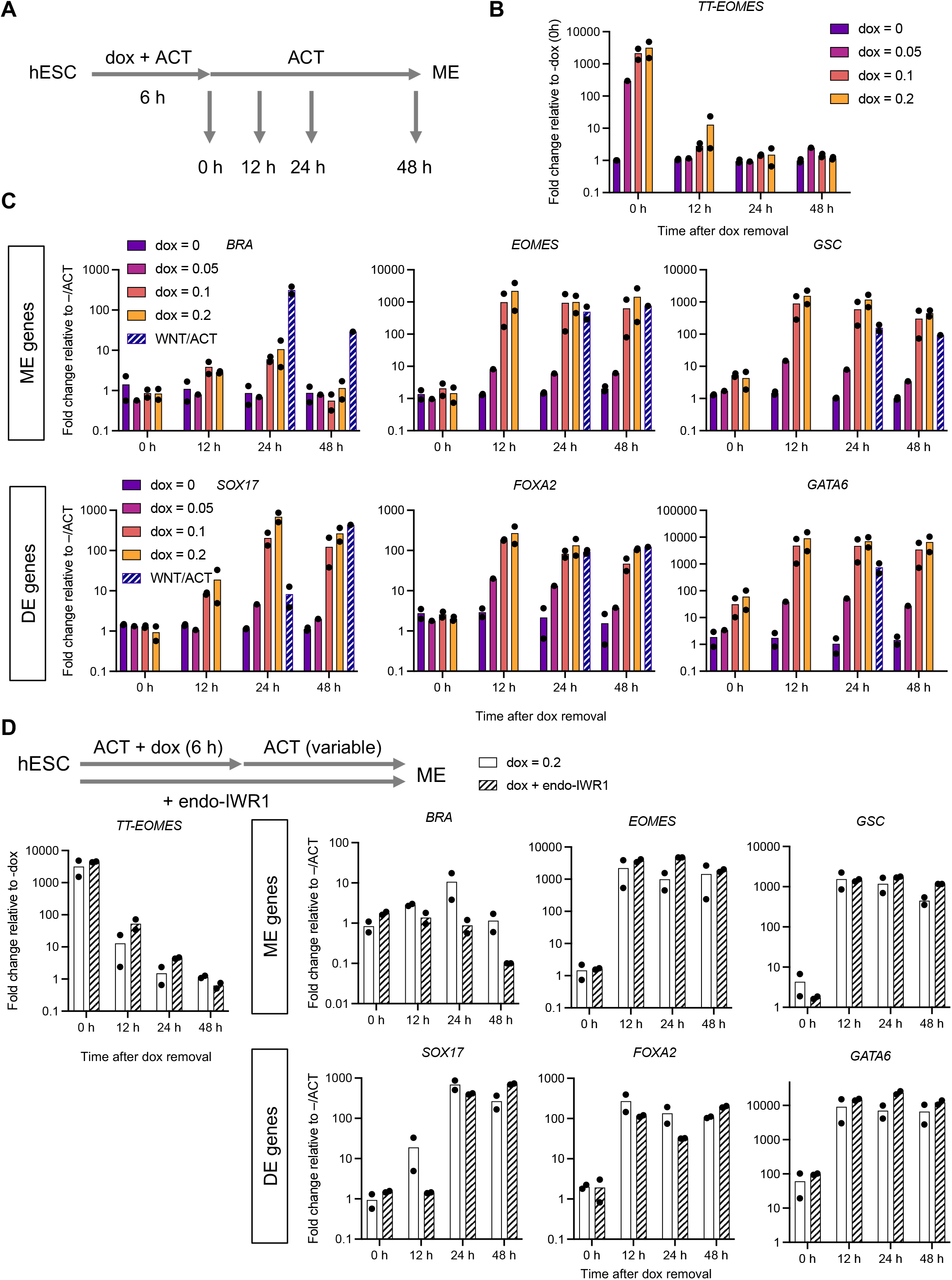
Exogenous EOMES expression bypasses WNT priming to induce ME differentiation. **A)** Protocol to assay ME differentiation in response to a pulse of TT-EOMES expression induced by doxycycline (dox). **B–C)** Time course analysis of (B) *TT-EOMES* and (C) endogenous ME gene expression following transient *TT-EOMES* induction (6 h with variable dox concentration). Cells were collected at 0, 12, 24, and 48 h during the Activin phase for RT-PCR measurements. Data points represent the mean fold change relative to pluripotency levels in the parental line (–/ACT, 24 h). The expression levels in the parental cell line treated with Wnt followed by ACT for 24 h or 48h are shown for comparison. Bars represent the mean across n = 2 biological replicates (black dots). **D)** Similar to (B) except that endo-IWR1 (1 μM) was added to block Wnt signaling. The bar plots show the mRNA levels of ME genes at different time points during the ACT phase with or without endo-IWR1.

Consistent with a previous study (Pfeiffer et al., 2018), we also found that EOMES can induce the expression of *WNT3* (Figure S8D). To eliminate the possibility that *TT-EOMES* drives hESC differentiation through the production of endogenous WNT, we repeated our experiments in the presence of endo-IWR1. When applied to the unmodified hESCs in the WNT/ACT protocol, endo-IWR1 completely blocks the activation of ME and DE genes (Figure 5B,C). In contrast, endo-IWR1 has little effect on the ME and DE gene expression driven by *TT-EOMES* (Figure 7D). These results strongly suggest that EOMES is an effector of Wnt priming, and exogenously expressed EOMES can drive ME differentiation in the absence of WNT signaling.

## Discussion

EOMES is a TF expressed during early development that is critical for endoderm and cardiac mesoderm formation in mouse and humans, as EOMES knockout completely abolishes these lineages (Arnold et al., 2008; Costello et al., 2011; Kartikasari et al., 2013; Li et al., 2019; Pfeiffer et al., 2018; Teo et al., 2011). It is also thought that EOMES is at the top of the ME gene regulatory network hierarchy, because it is one of the earliest genes that is induced during ME differentiation (Teo et al., 2011) and exogenous expression of EOMES in the presence of Activin promotes the expression of endodermal genes (Ameele et al., 2012). However, because EOMES engages in a complex gene network with WNT and Activin, e.g. EOMES can be activated with WNT signaling, and WNT3 is also a direct target of EOMES (Pfeiffer et al., 2018), it was not clear if EOMES could completely bypass WNT and directly induce ME differentiation with Activin. In fact, it was proposed that the activation of WNT signaling downstream of EOMES is critical for ME and cardiac differentiation (Pfeiffer et al., 2018). Here we show that induction of EOMES at near physiological levels in the presence of Activin is *sufficient* for ME and DE induction without the involvement of β-CATENIN signaling. While there is prior literature showing EOMES is a cofactor for SMAD2 in ME differentiation, we specifically found EOMES and SMAD2 co-binding enriched in genes sensitive to WNT priming (Fig. S6E), which establishes a critical link between WNT and Activin signaling in ME differentiation and provides a potential mechanism for WNT priming and memory (Yoney et al., 2018).

EOMES binds to its own promoter along with SMAD2 and together EOMES and SMAD2 drive ME differentiation (Kartikasari et al., 2013; Teo et al., 2011). One question raised by our study is then why this positive feedback loop is not stably activated with WNT priming alone, i.e. why does *EOMES* expression decrease when cells are returned to neutral media after WNT priming? In our protocol EOMES is activated by 24 h WNT (mRNA level is increased by ~200 fold in comparison to E7), but Activin can induce its expression further (~1500 fold in comparison to E7) (Fig. 1B). It is possible that EOMES is already associated with its own promoter in WNT but cannot drive itself to a high level without SMAD2. Alternatively, EOMES concentration may need to reach a critical level to jumpstart the feedback, and this level is not reached during the WNT priming phase. Further studies are needed to distinguish among these possibilities. Further investigation is also required to determine the mechanism by which EOMES and Activin/SMAD2 lock in the ME fate. One possibility is that EOMES suppresses pluripotency factors or alternative differentiation pathways (Teo et al., 2011; Tosic et al., 2019). Our proposed mechanism for WNT priming is summarized in Fig. 8.

**Figure 8:**
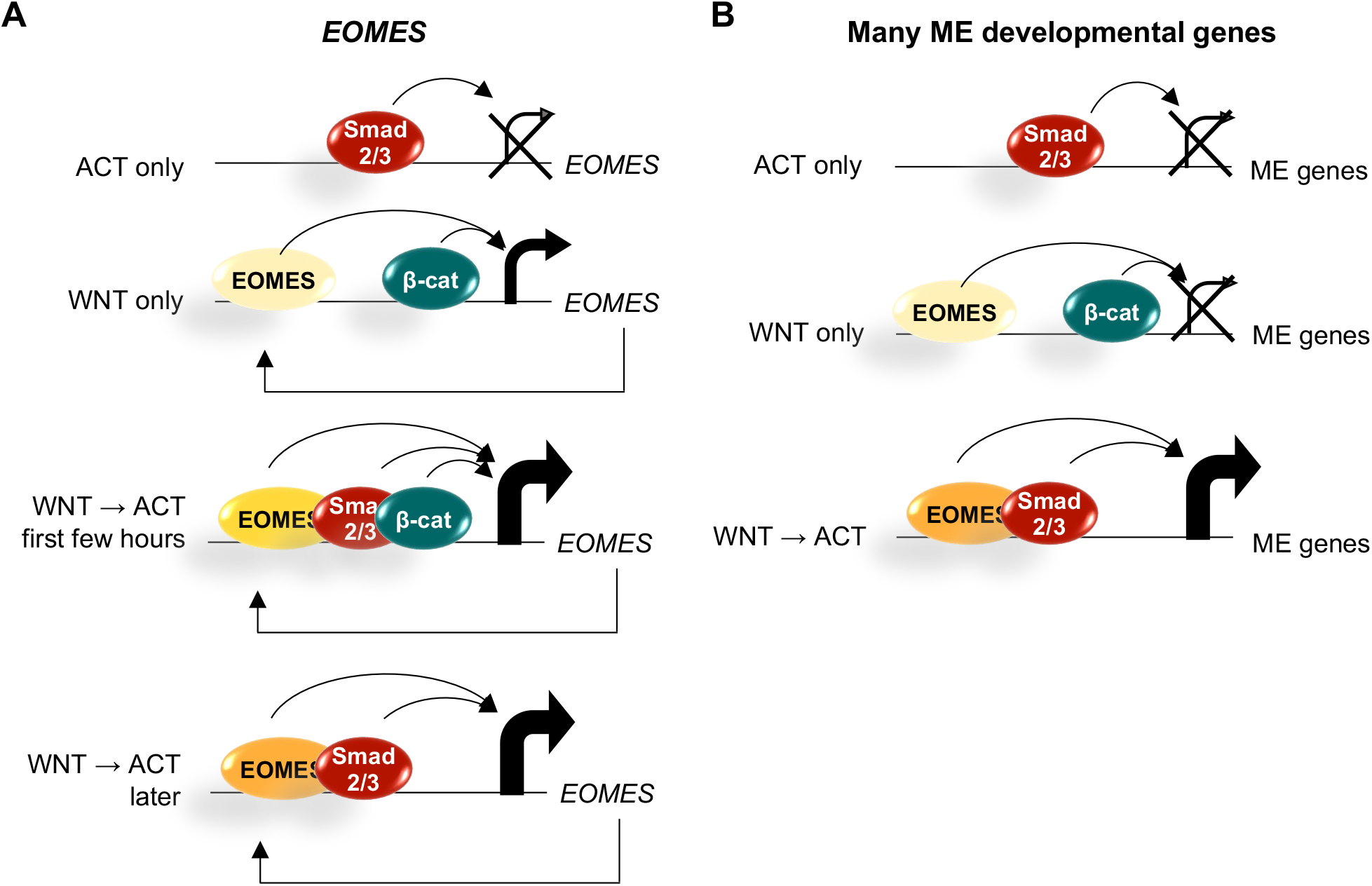
EOMES is the main effector of WNT priming. **A)** WNT signaling through β-CATENIN induces low levels of EOMES expression, which are further boosted by additional factors including SMAD2/3 and EOMES itself. When EOMES levels are sufficiently high, its expression can be maintained in the absence of WNT/β-CATENIN. **B)** Likewise, sufficiently high levels of EOMES together with SMAD2/3 can drive additional genes required for complete ME differentiation.

Chromatin modifications have long been implicated in cellular memory as the term epigenetics implies. In *Xenopus* a delay between WNT signaling and transcriptional onset was linked to an epigenetic mechanism in which β-CATENIN recruits a histone methyltransferase to target genes and is required for their activation at the mid-blastula transition (Blythe et al., 2010). The competence of the *Xenopus* blastula to respond to WNT signals was also shown to have a chromatin component (Esmaeili et al., 2020). Closer to our study, Tosic et al. showed that EOMES and BRA are required for ME gene expression in mouse and that their binding is associated with open chromatin. We did not find evidence for drastic changes in chromatin opening with WNT priming. β-CATENIN binding sites are already highly accessible in hESCs, which is likely because these sites are occupied by TCF/LEF proteins (Estarás et al., 2015), and β-CATENIN binding only mildly enhances the local ATAC-seq signals. In addition, the presence of β-CATENIN does not alter the accessibility near the SMAD2/3 sites, consistent with the idea that SMAD2/3 binding is not affected by WNT priming. Likewise, SMAD2/3 in the Activin alone condition does not enhance chromatin accessibility over the β-CATENIN binding sites. Taken together, these findings suggest that in contrast to what would be predicted from earlier models on ME differentiation, the chromatin opening and the direct binding of β-CATENIN and SMAD2/3 are independent events along the path to ME differentiation. Additionally, by superimposing our ATAC-seq analysis with our analysis of prior published data sets, we demonstrate a consistency among multiple similar protocols for mesendoderm induction.

Gene regulation by TGFβ signaling in different cell types often involves alternative SMAD2/3 binding partners. For example, SMAD2 binding in hESCs was proposed to depend on OCT4, NANOG, and FOXH1 (Attisano et al., 2001; Brown et al., 2011; Chen et al., 1996; Mullen et al., 2011; Teo et al., 2011), and on EOMES and GATA6 in definitive endoderm (Li et al., 2019). However, we find that SMAD2 binding does not seem to differ in pluripotent hESCs vs when compared to ME cells. This may be due to the fact that SMAD2/3 co-binders, OCT4, NANOG, and FOXH1, all have similar levels in these two cell types. Note that this is different from the situation in definitive endoderm cells where OCT4 and NANOG are significantly downregulated, which likely allows SMAD2 to target other loci. Therefore, we propose that the induction and subsequent binding of EOMES provides transcriptional activation downstream of SMAD2/3 binding, potentially by EOMES-dependent recruitment of chromatin modifiers (Kartikasari et al., 2013). Our findings reveal that one essential function of WNT signaling is the induction of EOMES, providing a simple mechanism of signaling integration that drives cellular differentiation. How induction of EOMES may impart a dose-dependent response to Activin/SMAD2 signaling that is responsible for further patterning the ME remains an open question.

## Materials and methods

### Human embryonic stem cell culture

Experiments were performed with the RUES2 hESC line (XX female; US National Institutes of Health, human ESC registry no. 0013), which was authenticated by STR profiling. Cells were routinely tested for mycoplasma contamination. For maintenance, hESCs were grown in HUESM medium that was conditioned by mouse embryonic fibroblasts and supplemented with 20 ng/mL bFGF (MEF-CM). Cells were grown on tissue culture dishes coated with Geltrex (Thermo Fisher Scientific, Waltham, MA) and passaged as aggregates using Gentle Cell Dissociation Reagent (STEMCELL Technologies, Vancouver, Canada). For experiments, single cells were obtained by dissociation with Accutase (STEMCELL Technologies) and seeded at low-density in optical-quality plastic tissue culture dishes (50k per dish) or 24-well plates (5k per well) (ibidi, Martinsried, Germany) in TeSR-E7 medium (STEMCELL Technologies) supplemented with 10 μM Rock inhibitor (Y-27632, Abcam, Cambridge, MA). Prior to seeding, dishes or plates were coated with 10 μg/mL Laminin-521 (BioLamina, Sundbyberg, Sweden) in PBS +Ca/+Mg for 2 hours at 37 °C or overnight at 4 °C. For the 2-day protocol in which cells were switched from Wnt3a to Activin A (R&D Systems, Minneapolis, MN), the samples were washed with PBS +/+ before adding fresh medium. 10 μM Rock inhibitor was maintained throughout the duration of the experiment. Other small molecules were used at the following concentrations and were replaced every 24 hours: 2.5 μM CHIR99021 (EMD Millipore), 1 μM IWP-2 (Stemgent), 1 μM endo-IWR-1 (Tocris), and 10 μM SB431542 (Stemgent).

### Immunofluorescence

Cells were rinsed once with PBS, fixed with 4% paraformaldehyde (Alfa Aesar, Thermo Fisher Scientific, Tewksbury, MA) for 20 minutes at room temperature, and then rinsed twice and stored in PBS. Cells were blocked and permeabilized with blocking buffer (2% bovine serum albumin and 0.1% Triton X-100 in PBS) for 30 minutes at room temperature. Cells were incubated with primary antibodies in blocking buffer overnight at 4 °C and then washed three times with 0.1% Tween-20 in PBS (PBST). The following primary antibodies and dilutions were used: SOX2 (rabbit monoclonal, Cell Signaling Cat. No. 3579, 1:200), BRACHYURY (goat polyclonal, R&D Systems AF2085, 1:150), BRACHYURY (rabbit monoclonal, R&D Systems MAB20851, 1:400), EOMES (mouse monoclonal, R&D Systems MAB6166, 1:400), GSC (goat polyclonal, R&D Systems AF4086, 1:200). Cells were incubated with secondary antibodies (diluted 1:1000): Alexa Fluor 488, 555, or 647-conjugated (Invitrogen Molecular Probes, Thermo Fisher Scientific) and DAPI nuclear stain in blocking buffer for 30 minutes at room temperature, and then washed twice with PBST and once with PBS.

### Imaging and analysis

Wide-field images were acquired on an Olympus IX-70 inverted microscope with a 10x/0.4 numerical aperture objective lens. Tiled image acquisition was used to acquire images of large areas in four channels corresponding to DAPI and Alexa Fluor 488, 555, and 647. Image analysis was performed using custom software in MATLAB as described in (Yoney et al., 2018). Nuclei segmentation and signal quantification were performed on background-corrected images as follows. The DAPI image was thresholded to generate a binary image separating the foreground (nuclei) from the background. The DAPI image was then filtered with a median and sphere filter with parameters matching the expected size of individual nuclei. Local maxima corresponding to individual nuclei were detected using the MATLAB extended-maxima transform function. Maxima falling within the foreground were used as seeds for watershed segmentation, which was also restricted to the foreground and was used to obtain a labeled object corresponding to each nucleus within the image. The results of the segmentation were used as a mask to obtain the median per cell nuclear intensity in each channel for 5-10k cells per condition from which histograms were generated.

### ATAC-sequencing

Chromatin accessibility profiling was carried out using the Omni-ATAC-sequencing protocol as an attempt to reduce mitochondrial DNA in our samples (Corces et al., 2017). We obtained single cells using Gentle Cell Dissociation Reagent (STEMCELL Technologies), rather than Accutase or other enzyme-based reagents, which we found disrupted downstream processing steps. Cells (100,000 per condition) were resuspended in 50 μL lysis buffer (0.1% NP-40, 0.1% Tween-20, 0.01% Digitonin) prepared in resuspension buffer (10 mM Tris-HCl pH 7.4, 10 mM NaCl, and 3 mM MgCl2 in water) for 3 minutes on ice. We optimized the lysis time so that the outer membrane was disrupted as indicated by Trypan blue staining of the nuclei but that the nuclei remained intact. The lysis buffer was removed by washing with 0.1% Tween-20 in resuspension buffer, and the transposition was subsequently carried out in 50 μL reaction volume containing 25 μL 2X TD buffer and 2.5 μL Tn5 transposase (available as individual products upon request from Illuminia). The transposition reaction was incubated at 37 °C for 30 minutes in a thermomixer set to 1,000 r.p.m. The transposed DNA was isolated using the Qiagen MinElute kit and was eluted in 30 μL of water in the final step. Library preparation was carried out using 10 μL of transposed DNA per sample as described previously with 10 - 11 cycles of amplification (Buenrostro et al., 2015). Our DNA tended to be under digested, so we performed doubled-sided bead purification using AMPure XP beads (Beckman). Libraries were sequenced as paired-end 75 bp reads, multiplexing all samples per experiment (7) on one lane of the Illumina High-Seq 500 platform at The Rockefeller University Genomics Resource Center. Two biological replicates for each condition were collected, processed, and sequenced from independent experiments.

### ChIP

For ChIP experiment in Fig. 2D, we used a SMAD2/3 antibody (R&D systems, AF3797) in 2X10^6 hESCs that were treated with Activin for 2 h or 24 h with or without Wnt priming. We used the Pierce magnetic ChIP kit and followed the procedure provided by the kit. Known SMAD2 binding sites near *GSC*, *NODAL*, and *CER1* genes were selected for subsequent qPCR analysis based on previously published SMAD2/3 ChIP-seq data (Kim et al., 2011; Tsankov et al., 2015). A region near *CER1* gene that has low SMAD2/3 ChIP-seq signal was chosen as the negative control. The primer sequences used for ChIP are included in Table S3.

### Bioinformatics analysis

RNAseq data were aligned by Rsubread, and the read count associated with genes were analyzed by featureCounts. The differential fold changes and effective p-values were obtained using DESeq2 (Love et al., 2014). The data in Fig. 1C include those genes that show a significant change (adjusted p-value < 0.01) in at least one of the four conditions relative to E7: (1) E7/ACT, (2) 24 h WNT, (3) WNT/E7, or (4) WNT/ACT. The k-mean clustering was done using MATLAB *kmeans* function. We used a number of clusters (8 in total) such that the final clusters do not change based on the randomized starting conditions for each run of the algorithm. For easier description, we combined three of them into cluster 3 (genes more activated by WNT/ACT than other three conditions), and another three into cluster 4 (repressed genes). We provided a list of genes belonging to each cluster (or sub-cluster) in supplementary Table S1.

The ChIP-seq and ATACseq data were aligned with bowtie2 for the hg19 reference genome and visualized using deepTools bamCoverage function. The peaks were identified using MACS2 narrow peak calling. The area underneath the peak (like in Fig. 2A, 3B, and 3D) were calculated using MultiCovBed function. The heatmaps in Fig. 3A and C were generated using deepTools computeMatrix and plotHeatmap. The differential peaks in Fig. 6B were identified using DESeq2 based on the areas underneath each peak (*P* < 0.01). In Fig. 6C, we collected the transcription start site (TSS) and transcription end site (TES) of genes in different categories and examined if there are any WNT/ACT enhanced ATAC-seq peaks that fall between 20kb upstream of the TSS and 20kb downstream the TES. The enriched motifs in the WNT/ACT enhanced ATAC-seq peaks were identified by AME (McLeay and Bailey, 2010) with the HOCOMOCOv10 database (Kulakovskiy et al., 2016) against other ATAC-seq peaks that are not sensitive to the WNT/ACT condition.

## Acknowledgements

We thank current and past members of our group for feedback on this work. We are especially grateful to Dr. Riccardo De Santis, Shu Li, and Peter Ingrassia for technical support in completing the revisions.

## Data availability

Published datasets used in this study include β-CATENIN ChIP data (GSM1579346), SMAD2/3 ChIP data from pluripotent hESCs (GSM727557), SMAD2/3 ChIP data from human mesendoderm cells differentiated from hESCs (GSM1505750), and EOMES ChIP data (GSM1505630 and GSM1505631). The RNA-seq and ATAC-seq data generated from this study have been deposited in NCBI’s Gene Expression Omnibus.

