## Supplementary information for "Mechanisms Underlying WNT-mediated Priming of Human Embryonic Stem Cells"

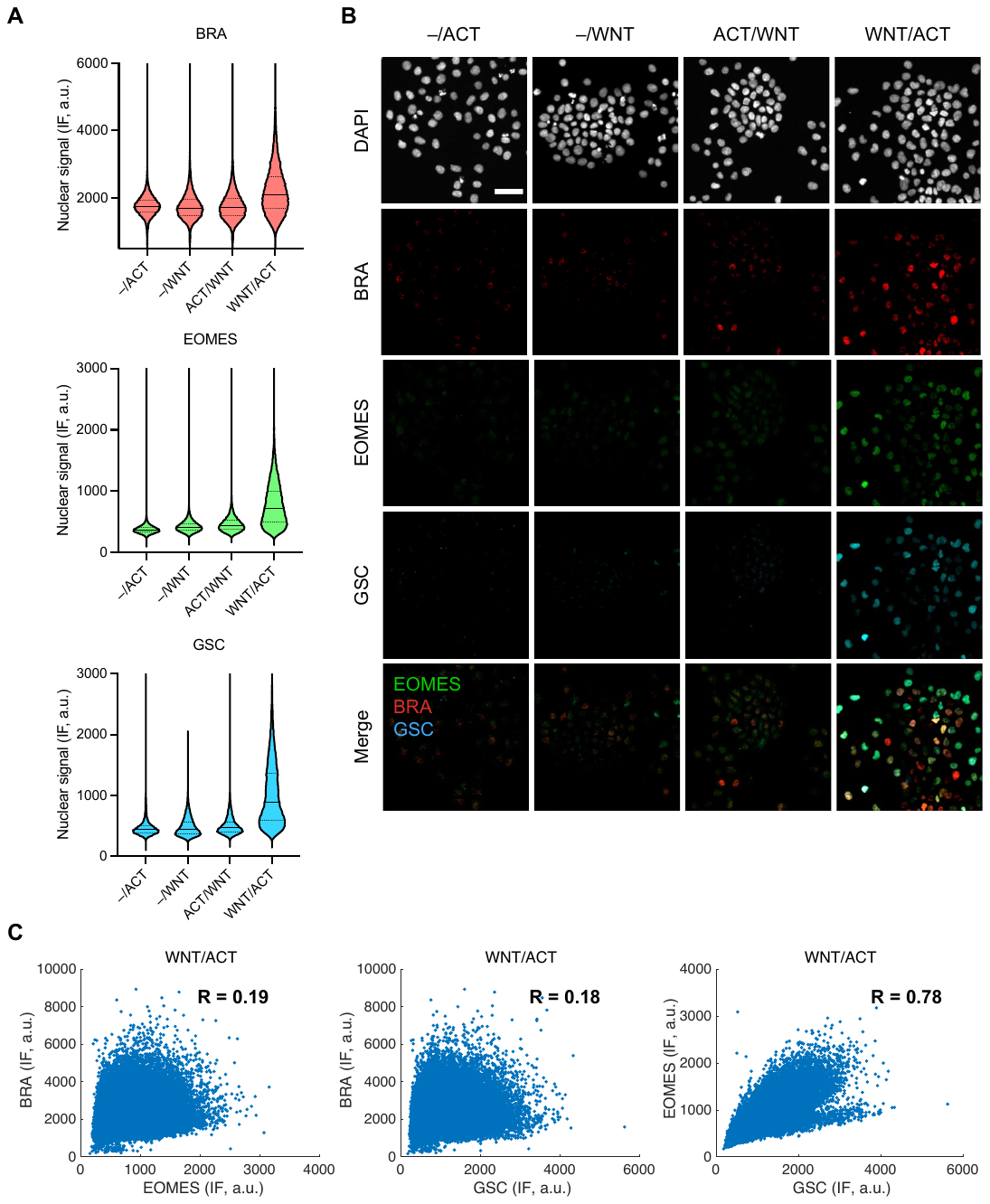

**Figure S1: Reverse priming does not lead to ME differentiation. A)** ME differentiation requires Wnt (100 ng/mL, 24 h) priming prior to 10 ng/mL Activin treatment. The expression of the three ME genes was measured after WNT/ACT or ACT/WNT protocols and with WNT alone on the second day (–/WNT). Only the WNT/ACT protocol shows expression above background signals quantified in pluripotency conditions (–/ACT). Violin plots of the median nuclear immunofluorescence (IF) signals of three ME markers (EOMES, BRA, and GSC) quantified in single cells (n > 20,000 cells per condition). Solid line, median; Dashed lines, upper and lower quartiles. **B)** Example images corresponding to the analysis shown in (A). Scale bar, 50 µm. **C)** Scatter plots of the data shown in B from the WNT/ACT condition. EOMES and GSC are more highly correlated with each other than with BRA. R, Pearson correlation coefficient.

**Figure S2: Wnt priming is achieved with CHIR. A)** WNT signaling was activated by 24h CHIR99021 (CHIR, 2.5 µM) treatment, and its effect on ME differentiation ± 10 ng/mL Activin was analyzed by immunofluorescence. Violin plots of the median nuclear immunofluorescence (IF) signals of three ME markers (EOMES, BRA, and GSC) quantified in single cells (n > 10,000 cells per condition). Solid line, median; Dashed lines, upper and lower quartiles. **B)** Example images corresponding to the analysis shown in (A). Scale bar, 50 µm. **C)** Scatter plots of the data shown in B from the ME conditions (WNT/ACT and CHIR/ACT). Similar to the data shown in Figure S1, EOMES and GSC are more highly correlated with each other than with BRA. R, Pearson correlation coefficient.

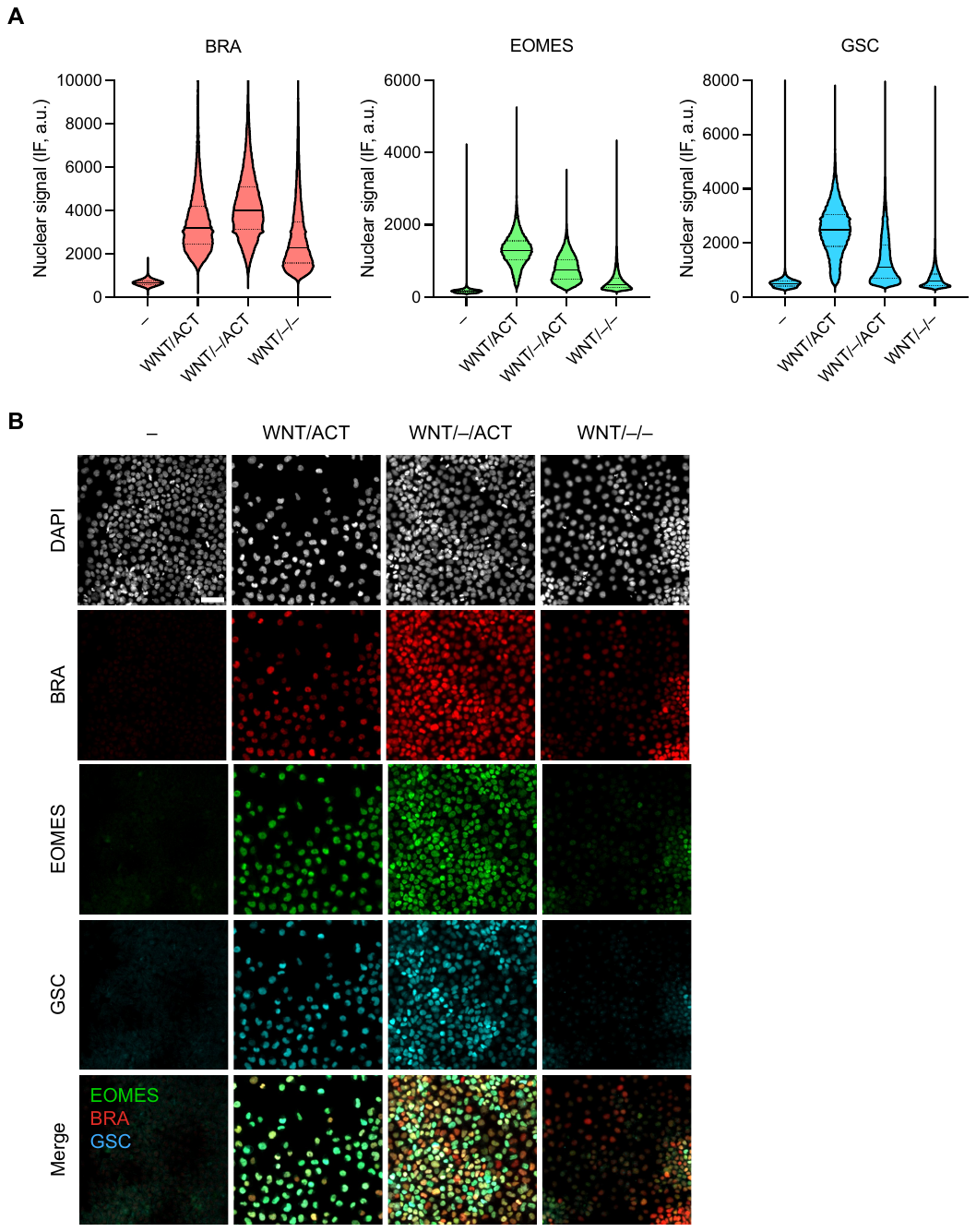

**Figure S3: Additional data supporting the Wnt memory phenomenon.** Independent repeat of the conditions shown in Figure 1D, E in which SB was not included. Violin plots of the median nuclear immunofluorescence (IF) signal for three ME markers (BRA, EOMES, and GSC) quantified in single cells (n > 40,000 cells per condition). Solid line, median; Dashed lines, upper and lower quartiles. **B)** Example images corresponding to the analysis shown in (A). Scale bar, 50 µm.

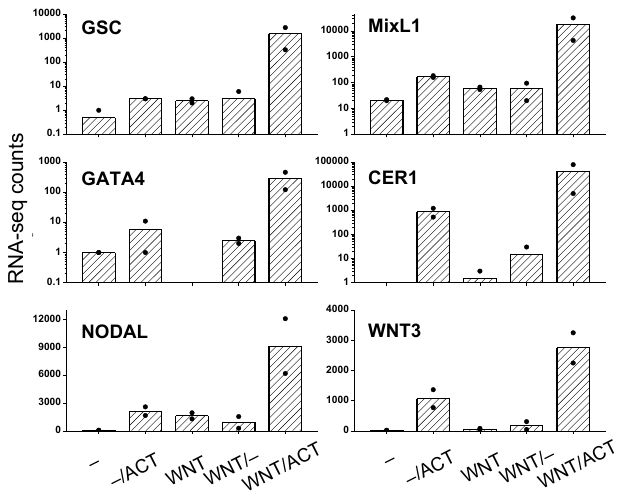

**Figure S4: mRNA Expression levels of the genes shown in Figure 2A.** RNA counts were extracted from RNA-seq measurements (two biological replicates for each condition). All genes show significantly higher expression in the WNT/ACT condition relative to all other conditions.

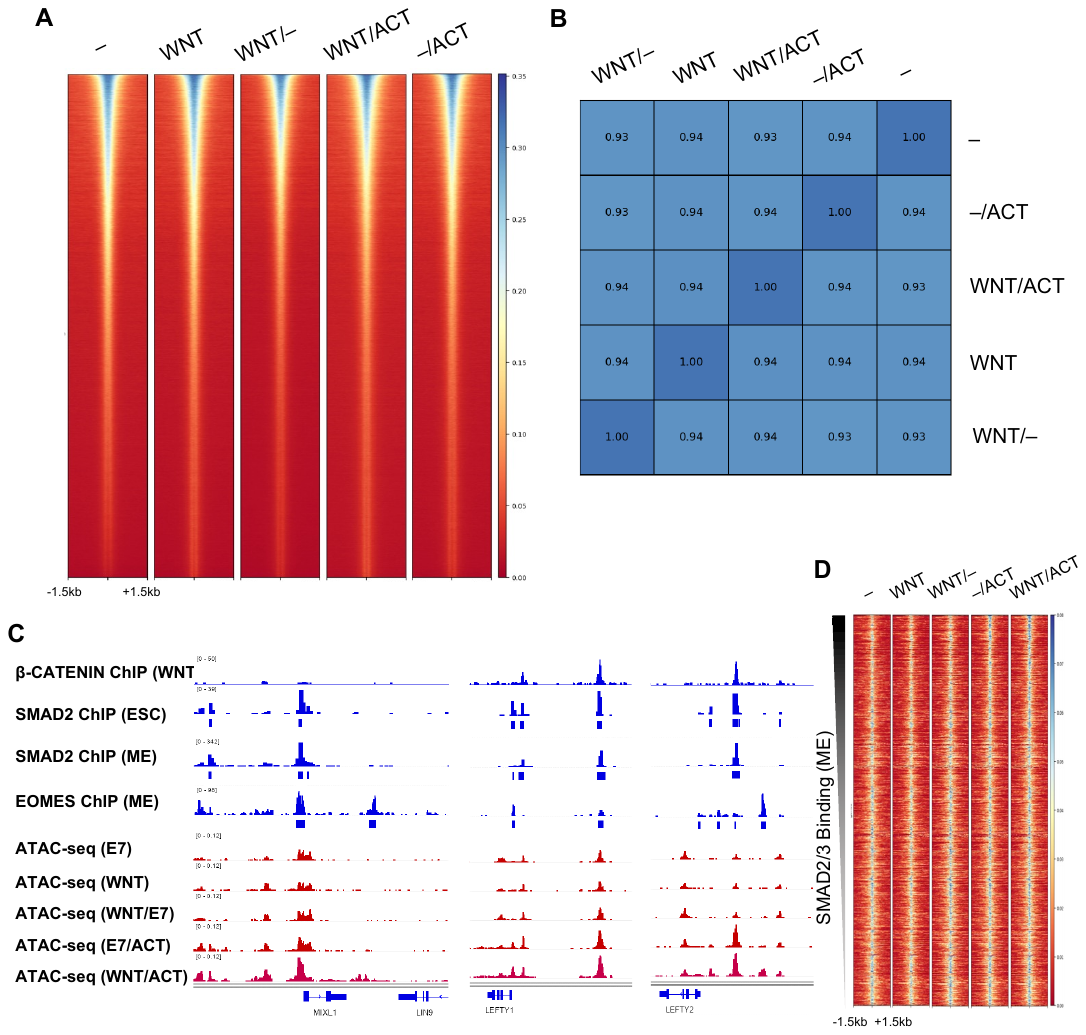

**Figure S5: ATAC-seq signals remain largely unchanged after WNT and/or Activin treatment. A)** Heatmap of ATAC-seq peaks measured in five conditions as indicated at the top. The peak regions called in both replicates from each condition were combined and sorted in descending order based on their intensities in the E7 condition (–). All five heatmaps were plotted using the same sorted regions. The similarity of these plots indicates that there is no global rearrangement of open chromatin regions. **B)** Spearman rank-order correlations among the five ATAC-seq datasets are all > 0.93. **C)** β-CATENIN, SMAD2, and EOMES ChIP-seq, as well as ATAC-seq data near *MIXL1* and *GATA4* genes. The enhanced ATAC-seq peaks in WNT/ACT overlap with the ChIP-seq peaks of some of these factors. **D)** Same as in Figure 3D except that that SMAD2/3 binding peaks were extracted from the ChIP-seq data in ME cells.

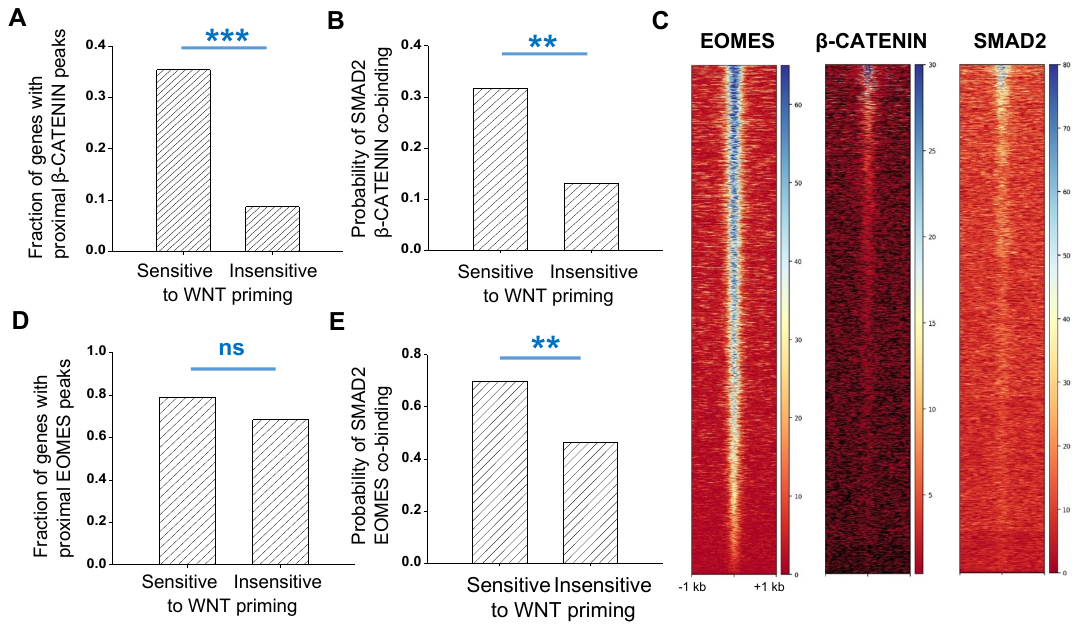

**Figure S6: Binding of β-CATENIN and EOMES to WNT-primed genes. A)** Fraction of genes that are proximal to β-Catenin ChIP-seq peaks, i.e., at least one β-Catenin ChIP-seq peak can be found over within TSS ± 5kb. This fraction is compared among genes that are Wnt-primed, and those that are not Wnt-primed (*P<0.05, **P< 0.01, ***P<0.001, ns: not significant). **B)** Fraction of SMAD2/3 sites that co-localize with β-Catenin sites. For each gene, within the range of TSS ± 5kb, we looked for SMAD2/3 and β-Catenin ChIP peaks that are within 500 bp (co-localized peaks). The probability is calculated as the number of genes that contain co-localized peaks divided by the number of genes that contain SMAD2/3 peaks. **C)** Sorted ChIP-seq signals of β-CATENIN, SMAD2, and EOMES at the WNT/ACT enhanced ATAC-seq peaks. Most of these sites bind to EOMES, but not β-Catenin or SMAD2. **D–E)** Same as in (A) and (B) except for EOMES.

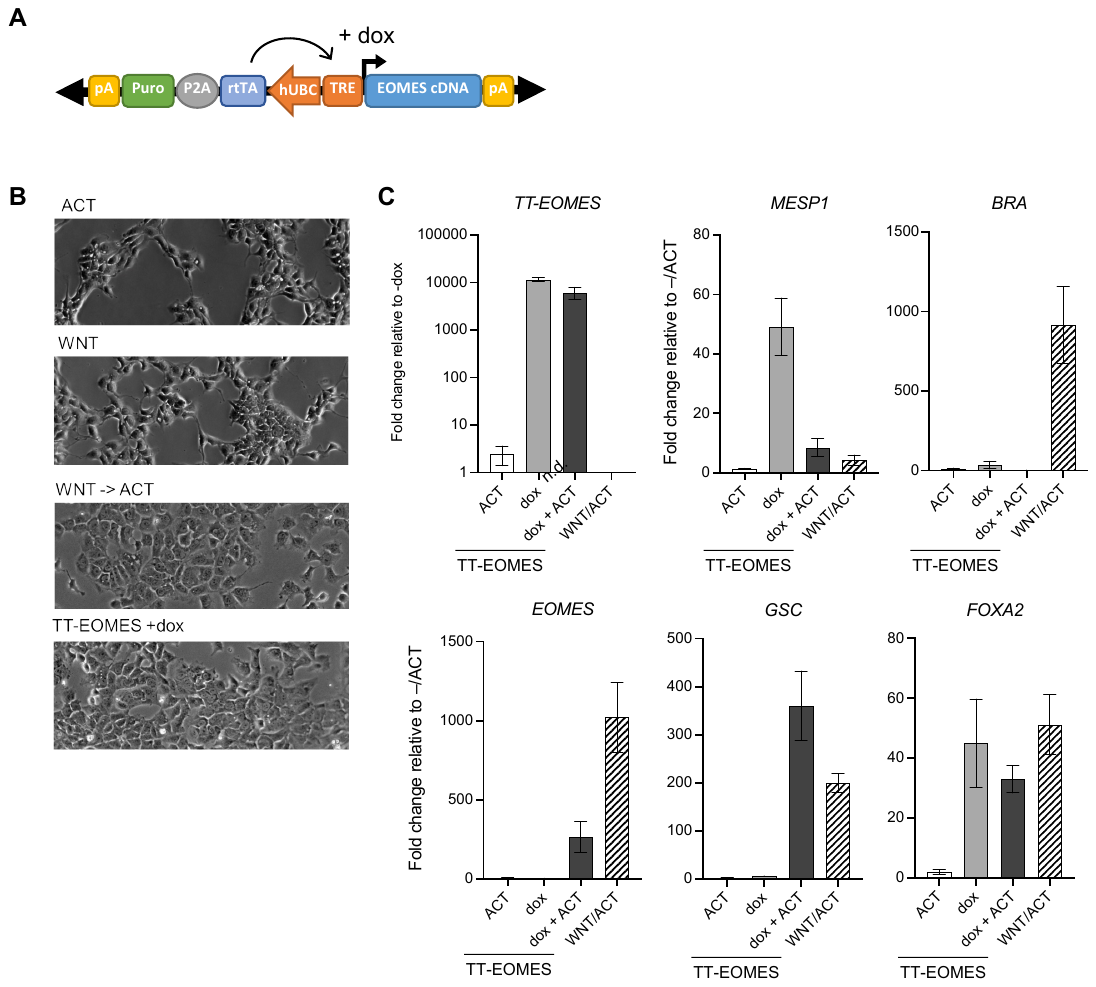

**Figure S7: Generation of a dox-inducible EOMES hESC line. A)** Construct integrated into hESCs to induce exogenous EOMES expression with doxycycline (dox). **B)** Phase contrast images showing that dox treatment of the TT-EOMES line induces morphological changes that are similar to those observed with WNT/ACT treatment of the unmodified parental line. Cells retain a pluripotent morphology with WNT or Activin only. **C)** TT-EOMES hESCs were treated with 24 h Activin (10 ng/mL), 48 h dox (1 µg/mL), or 48 h dox (1 µg/mL) + Activin (10 ng/mL). Expression levels after standard WNT/ACT treatment of the unmodified parental line are shown for comparison. Expression in each sample was normalized to GAPDH and then to the pluripotency levels in the parental line (–/ACT) or in the case of exogenous EOMES expression (*TT-EOMES*), to the levels in the absence of dox. Similar to what was reported previously, EOMES expression induces the marker of cardiac mesoderm, *MESP1*, and Activin treatment reduces *MESP1* expression. Error bars represent the standard deviation over technical replicates. n.d., not detected.

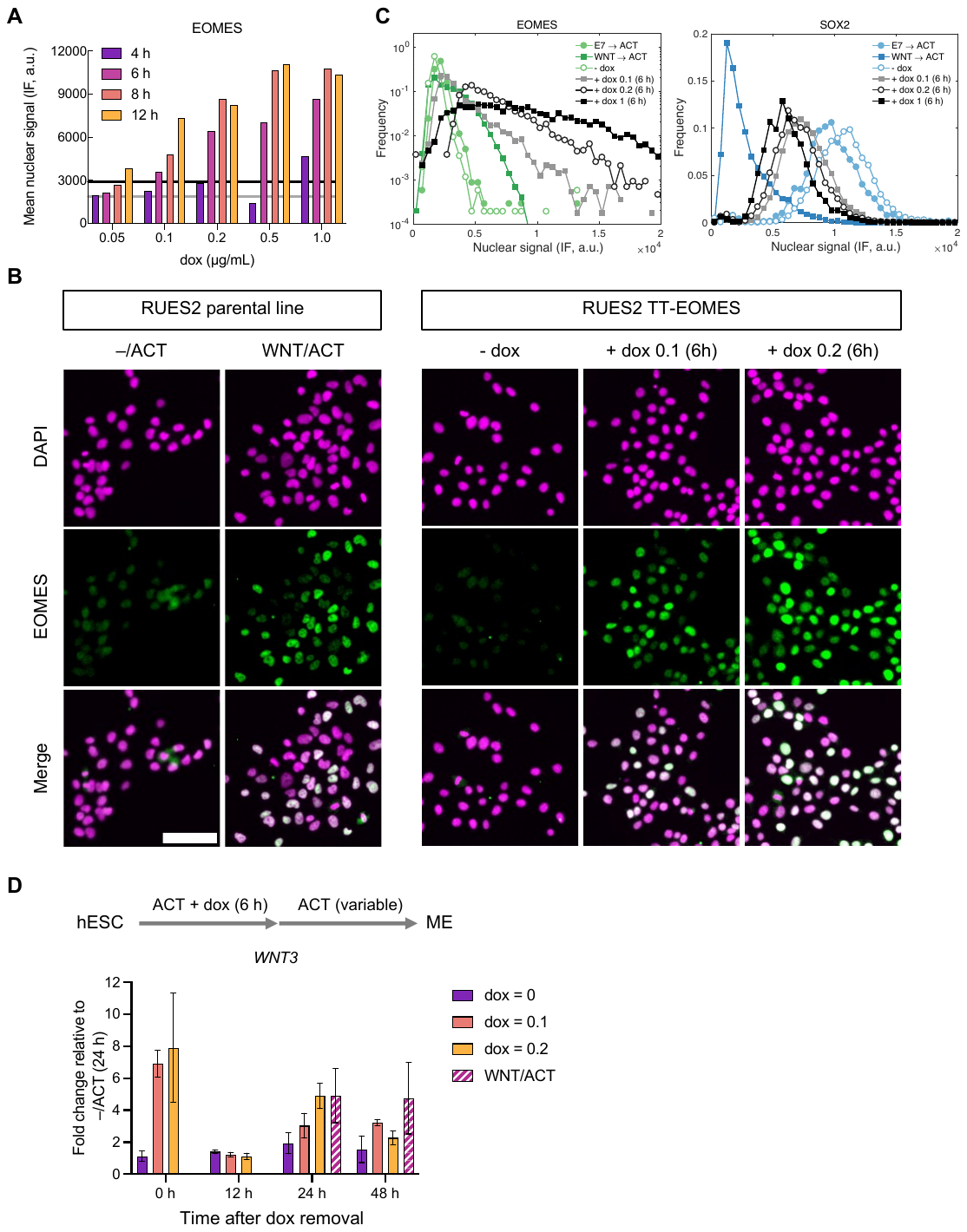

**Figure S8: Titration of TT-EOMES expression to achieve endogenous protein levels. A)** EOMES expression in a clonal TT-EOMES (isoform 1) hESC line induced with different dox levels plotted along the x-axis. Expression was analyzed by immunofluorescence at different time points after dox treatment with an EOMES antibody that recognizes both the exogenous and endogenous protein. The bars show the mean expression level per cell. The mean expression in the unmodified parental line after WNT/ACT treatment (black line) and the modified line in the absence of dox (gray line) are shown for reference. The median nuclear signal was quantified in n > 2000 signal cells per condition. **B)** Example images corresponding to the analysis shown in (B) for –/ACT, WNT/ACT, and 6h dox 0.1 and 0.2 µg/mL. Scale bar, 100 µm. **C)** Histograms corresponding to the data in (A) and (B) at 6h dox treatment along with the corresponding SOX2 data. **D)** *WNT3* mRNA expression for the same experimental procedure in Figure 7A. Cells were collected at 0, 12, 24, and 48h during the Activin phase for RT-PCR measurements. Data represents the mean fold change relative to pluripotency levels in the parental line (–/ACT). Error bars represent the standard deviation over technical replicates.

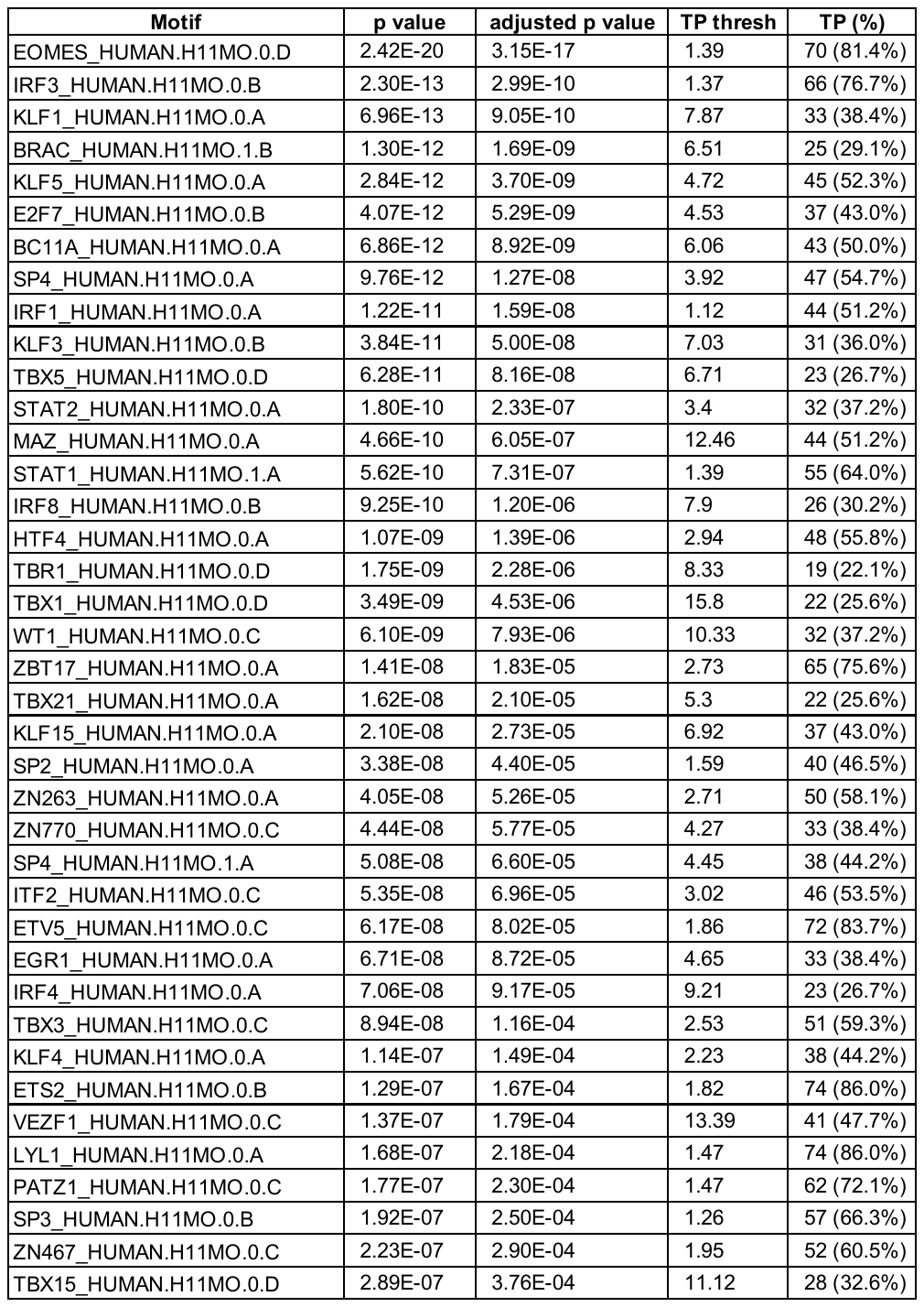

**Table S2. AME results for WNT/ACT enhanced peaks**

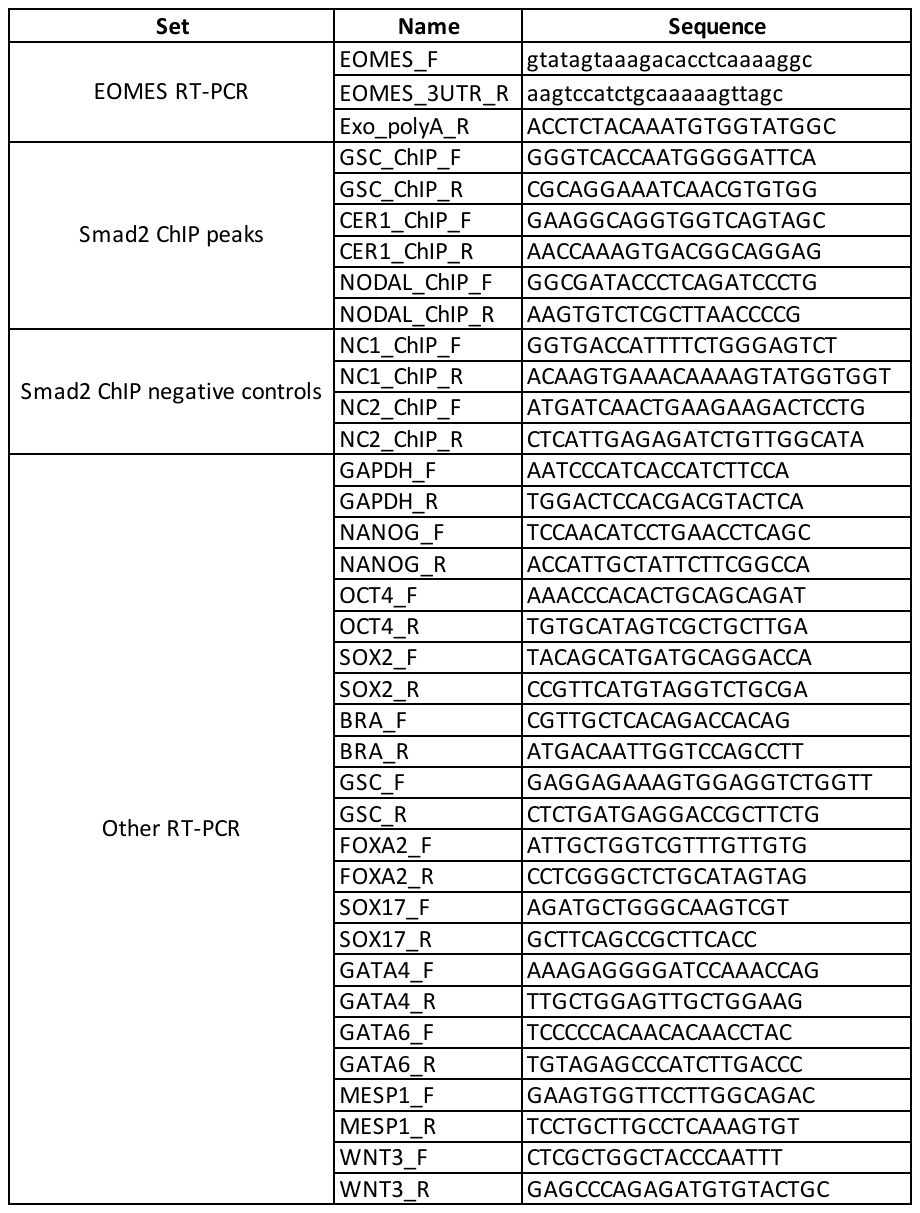

**Table S3. Primer Sequences**
